# The energy landscape for R-loop formation by the CRISPR-Cas Cascade complex

**DOI:** 10.1101/2023.03.17.533087

**Authors:** Dominik J. Kauert, Julene Madariaga-Marcos, Marius Rutkauskas, Alexander Wulfken, Inga Songailiene, Tomas Sinkunas, Virginijus Siksnys, Ralf Seidel

## Abstract

The discovery^1,2^ and the pioneering applications^3^ of CRISPR-Cas effector complexes have provided powerful gene-editing tools. The effector complexes are guided to the targeted genomic locus by the complementarity of their CRISPR RNA (crRNA)^4,5^. Recognition of double-stranded DNA targets proceeds via DNA unwinding and base-pairing between crRNA and the DNA target strand resulting in the formation of an R-loop structure^5,6^. Full R-loop formation is the prerequisite for the subsequent DNA cleavage. While the CRISPR-Cas technology is easy to use, efficient and highly versatile, therapeutic applications are hampered by the off-target effects due to the recognition of unintended sequences with multiple mismatches^7^. This process is still poorly understood on a mechanistic level^8,9^. Particularly, the lack of insight into the energetics and dynamics of the R-loop formation hinders a direct modelling of the R-loop formation for off-target prediction.

Here we set up ultrafast DNA unwinding experiments based on plasmonic DNA nanorotors to follow the R-loop formation by the Cascade effector complex in real time, close to base pair resolution. We directly resolve a weak global downhill bias of the energy landscape of the forming R-loop followed by a steep uphill bias for the final base pairs. We furthermore show a modulation of the landscape by base flips and mismatches. These data provide that Cascade-mediated R-loop formation occurs on short time scales in single base pair steps of sub-millisecond duration, but on longer time scales in six–base pair intermediate steps in agreement with the structural periodicity of the crRNA-DNA hybrid. We expect that the knowledge about the energy landscapes of R-loop formation of CRISPR-Cas effector complexes will pave the way for a detailed understanding and prediction of off-target recognition^10^.

---

DNA targeting by CRISPR-Cas effector complexes of Type I (Cascade), Type II (Cas9) and Type V (Cas12) is initiated by the crRNA-independent recognition of a short protospacer adjacent motif (PAM) on the DNA duplex. PAM recognition primes pairing between the PAM-proximal nucleotide (nt) stretch and the crRNA and is followed by further expansion of the R-loop structure^11^. R-loop formation up to the PAM-distal end of the target DNA triggers a conformational change of the effector complex that locks the R-loop in a stable conformation (Fig. 1a)^6,11–13^, which is a prerequisite for the subsequent DNA degradation. It is generally thought that R-loop expansion proceeds as a random-walk using fast and reversible single base pair steps such that whole genomes can be scanned within short time scales^14–16^. A proof of this diffusive nature of R-loop formation as well as important parameters of this process have so far remained unexplored, such as the free energy bias during R-loop expansion (downhill or uphill), details of the energy landscape leading to transient kinetic intermediates as well as the single base pair dynamics (Fig. 1b).

**Fig. 1.**
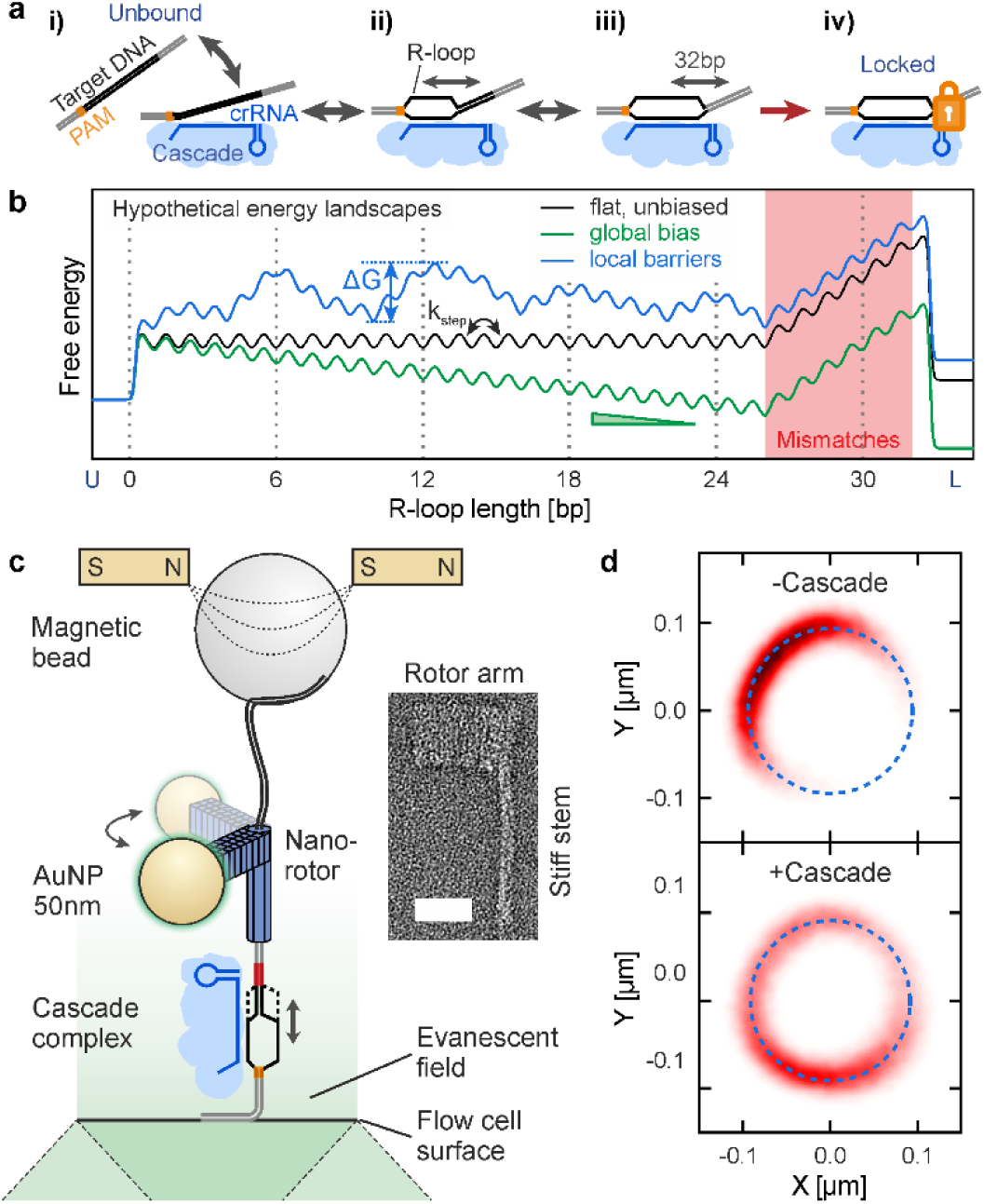
Single-molecule nanorotor measurements of R-loop formation. **a,** Scheme for R-loop formation by the Cascade complex (light blue with crRNA in blue): (i) PAM-mediated binding to the dsDNA target (grey), (ii, iii) R-loop priming and reversible expansion from the PAM and (iv) locking of the R-loop once it is fully formed. **b,** Different hypothetical free energy landscapes (coloured lines) for R-loop formation extending from the unbound (U) towards the locked (L) state. Mismatches (in red shaded area) represent energetic penalties and locally increase the energy landscape. **c,** Scheme of the nanorotor configuration. The DNA sequence of interest is attached on its bottom end to the surface of the fluidic cell and on its top end to the nanorotor consisting of a DNA origami nanostructure with a 56 nm long rotor arm at which a 50 nm gold nanoparticle (AuNP) is attached. A pair of magnets allows to stretch and twist the nanorotor construct via a magnetic bead that is attached at the top end of the origami structure through a 7.5-kbp DNA spacer. Imaging the light that is backscattered from the AuNP allows detection of the nanorotor rotations and thus monitoring the DNA untwisting during R-loop formation. A TEM image of the DNA origami nanostructure is shown on the right (scale bar: 25 nm). **d,** Projections of the nanorotor movement in the rotation plane before (top) and after (bottom) adding Cascade. The movement follows a circular path (dashed blue line).

Addressing these questions requires to follow R-loop formation at high spatio-temporal resolution for long observation times^16^. Since R-loop formation is coupled to extensive DNA untwisting of about 0.6 rad (34°) per bp, we set up ultrafast DNA twist measurements based on plasmonic DNA origami^17,18^ nanorotors. The L-shaped nanorotor structure consisted of a torsionally rigid six-helix-bundle stem and a 55 nm long rotor arm. The latter carried a 50 nm gold nanoparticle (AuNP) at its terminus for detecting the angular position of the nanorotor over long durations by light scattering^19^ (Fig. 1c, Extended Data Figs. 1,2). The rotor arm allowed the usage of comparably small AuNPs for optimum spatio-temporal resolution (see Supplementary Discussions S1-3) while maintaining a sufficiently large distance of the AuNP centre from the DNA for angular tracking.

The bottom end of the nanorotor was connected to a 47-bp DNA duplex carrying the desired target sequence which was furthermore rigidly attached to the surface of the fluidic cell. To maintain a vertical nanorotor orientation with minimal lateral fluctuations, we attached a 1 µm magnetic bead (Extended Data Fig. 3) via a 2.5 µm dsDNA spacer to the top end of the nanorotor stem and stretched it using magnetic tweezers. The setup supported molecule pulling and twisting as well as real-time measurement of the DNA length at 1000 fps using microscopy-based particle tracking^20^. Simultaneously, we illuminated the bottom glass slide through the objective in total internal reflection geometry with a 532 nm laser such that the resulting evanescent field of ∼200 nm depth was specifically illuminating the nanorotor (Fig. 1c, Extended Data Fig 4). The back-scattered light from exciting the surface plasmon of the AuNP was imaged at 3947 fps (see Supplementary Information for example video files) and the NP positions were tracked at 2.7 nm resolution (Extended Data Fig 5). For nanorotors with a torsionally constrained target sequence, the AuNP positions were located on a circular arc with a fixed mean and standard deviation of the angular position (Fig. 1d). Analysis of the angular fluctuations provided the torsional stiffness and the response time of this nanomechanical system yielding a spatio-temporal resolution for DNA untwisting of 3 bp at a time scale of ∼2 ms (see Supplementary Discussions S2 and Extended Data Fig 6). Compared to previous fast twist measurements^21,22^ (see Extended Data Table 1), our AuNP nanorotor uniquely provided a high spatio-temporal resolution in DNA untwisting measurements being supported during the required observation times in the hour range.

The integration of the nanorotors into magnetic tweezers also supported direct and fast torque measurements during DNA twisting of molecules for which the dsDNA linker part above the nanorotor was torsionally constrained (Extended Data Fig. 7). We were able to determine the torsional stiffness of DNA (torsional persistence length of *p_tor_* = 106 nm) as well as the torque during structural transitions of the twisted DNA, such as DNA denaturation^23^, DNA buckling^24^ and the formation of overwound p-DNA^25^ at a resolution of 1 pN nm within 38 ms.

We used the established nanorotor assay to study the dynamics of R-loop formation by the Type IE Cascade effector complex from *S. thermophilus* (Cascade)^2^ - a multimeric effector complex of five different Cas genes with a stoichiometry of Cse1_1_Cse2_2_Cas7_6_Cas5_1_Cas6e_1_^26^. It recognizes an AAN PAM and base pairs with the target sequence over a length of up to 32 bp. In order to allow an unconstrained R-loop expansion and shortening dynamics, we initially used a modified target sequence (further called T6) containing 6 mismatches to the crRNA at the PAM-distal end (Fig. 2a, top), which were shown to prevent locking over long durations^6,13^. We selected a nanorotor construct that had only a torsionally constrained target sequence (i.e. a nick within the linker region) to avoid any perturbation of R-loop formation by external torque. When adding Cascade into the flow cell, large counter-clockwise rotations of the nanorotors were observed indicating pronounced DNA untwisting due to R-loop formation (Figs. 1d, 2a). During an R-loop formation event the predominant untwisting angle was ∼15 rad, corresponding to a length of 26 bp in agreement with the number of complementary bases between DNA target and crRNA. The R-loop length was however constantly fluctuating towards shorter lengths indicating a dynamic and unconstrained R-loop (Fig. 2a, black section). Dynamic R-loops were frequently collapsing, such that multiple repetitive R-loop formation events were often observed (Extended Data Fig. 8). On a time scale of ∼60 min, R-loops could suddenly become stable at a length of 26 bp (Fig. 2a, red section), indicating R-loop locking despite of the 6 introduced mismatches.

**Fig. 2.**
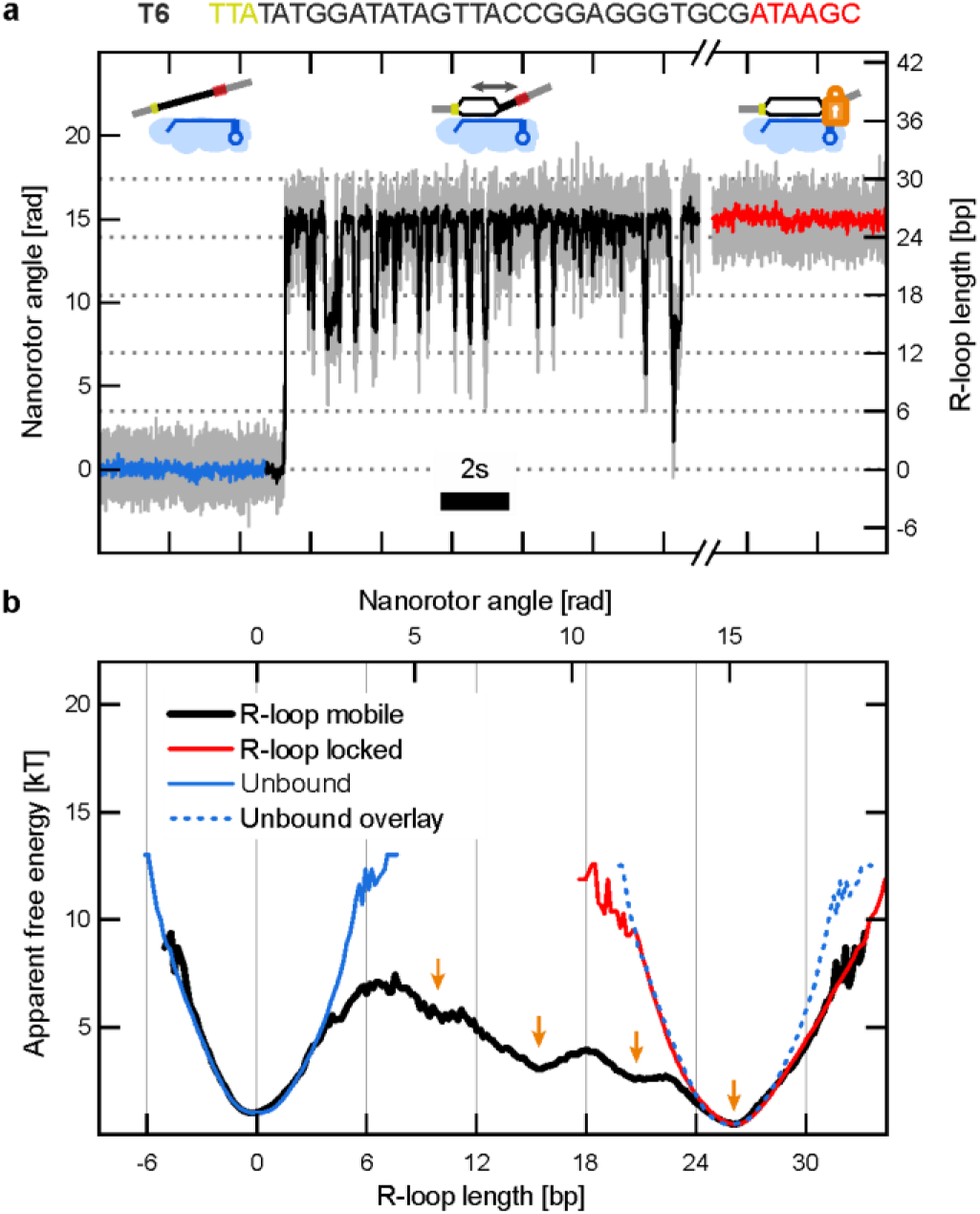
Real-time dynamics of R-loop expansion and shortening in the Cascade complex. **a,** Time trajectory of the angular position of the nanorotor (grey: raw data taken at 3947 fps; other colours: after 100-point sliding average). The target strand sequence is shown on top with PAM bases in yellow and PAM-distal mismatches with the crRNA in red, which strongly attenuate full R-loop zipping and locking. Upon Cascade binding, the initially constrained nanorotor (blue) started to diffuse within a large angular range (black) due to dynamic expansion and shortening of the unlocked R-loop. Finally, locking constrained the R-loop at a length of ∼26 bp (red). **b,** Apparent free energy landscapes (blue, black, red solid lines) calculated from the different sections of the trajectory in **a**. The landscapes are shifted along the energy axis for better comparison. Orange arrows indicate local free energy minima within the landscape of the unlocked R-loop. The blue dotted line is a shift of the landscape of the unbound state to the landscape of the locked state to allow comparison.

From the recorded DNA untwisting, we determined the occupancies of the different R-loop lengths and calculated the apparent free energy landscape for the R-loop formation (Fig. 2b, black line) using the Boltzmann distribution (see Supplementary Discussions S5). Compared to the real landscape, features in the apparent landscape are broadened (convolved) by the harmonic potential, which constrains the nanorotor fluctuations in absence of Cascade (Fig. 2b, blue line). The apparent energy landscape of the R-loop possesses a pronounced minimum at 0 bp indicative of the unbound state and noticeably a weak and approximately constant downhill bias up to 26 bp followed by a steep uphill bias.

This shows that R-loop formation for Cascade is energetically slightly preferential even in the unlocked state (∼5 *k_B_T* for the full 26-bp R-loop). On top of the global bias, the energy landscape exhibited a number of local energy minima and maxima possessing approximately a 6-bp periodicity (see arrows in Fig. 2b and occupancies in Extended Data Fig. 8a). This correlates well with the structure of the crRNA-target DNA duplex, which contains successive 5-bp duplex segments separated by 1 bp, where both the crRNA and the DNA bases are flipped out (Fig. 3a, top) allowing for a global parallel arrangement of the crRNA and the target strand (Fig. 3e). Flipped-base positions corresponded approximately to local energy maxima, while the centres of the 5-bp segments corresponded to energy minima. It has been suggested for the Type IE Cascade complex from *E.coli* that Lysin-rich “K-vise” motifs along the Cas7 subunits guide the PAM-distal dsDNA that is not yet incorporated in the R-loop^11^. dsDNA binding by these motifs reoccurring every 6 bp of the R-loop may be predominantly responsible for the energy minima along the measured energy landscape. To investigate whether the 6-bp periodicity was a general feature of the R-loop energy landscape of Cascade and to study the impact of different DNA bases at the flip positions, we obtained apparent energy landscapes for target variants in which the flipped DNA bases at positions 6, 12, 18 and 24 from the PAM where either changed to guanine (T6-Gflip target) or to thymine (T6-Tflip target, see Extended Data Table 2). For the three target variants the apparent energy landscapes where highly similar, exhibiting an approximately constant negative bias and local energy wells every 6 bp with the most pronounced minima located at the 3^rd^ and the 5^th^ 5-bp segment (Fig. 3a). The overall bias appeared to be similar for the T6-Tflip target while it was reduced for the T6-Gflip target indicating that the energy difference between flipped base stabilization by Cascade and disruption of the corresponding base pair in the target duplex is higher for GC base pairs.

**Fig. 3.**
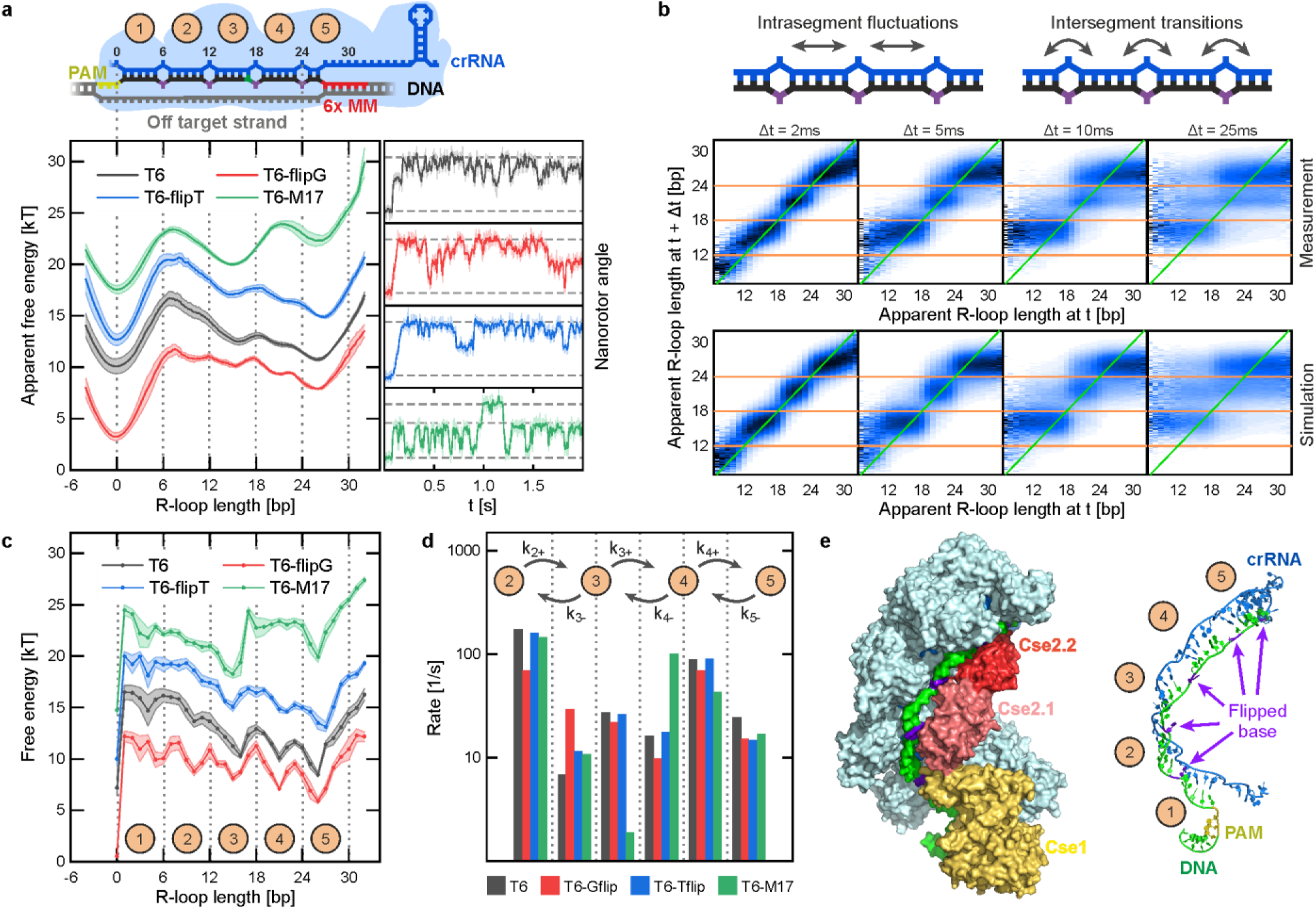
Long-range R-loop dynamics is governed by the 6-bp periodicity of the crRNA-target strand hybrid. **a,** Apparent free energy landscapes and example trajectories for the T6 target, its flipped base variants T6-Tflip and T6-Gflip as well as the T6 target with a C:A mismatch at position 17 (T6-M17). The R-loop scheme at the top is aligned with the R-loop length axis in the plots below to reveal the flipped base positions every 6 bp (dashed lines). For each construct the mean apparent free energies from five different molecules are shown with the shaded area indicating the s.e.m. Trajectories were recorded at 3947 fps (light colours) and smoothed with a 100-point sliding average (dark colours). Dashed lines indicate the R-loop extensions of 0 bp and 26 bp, as well as 17 bp for the T6-M17 target. **b,** Transition plots for R-loop length fluctuations on a T6 target indicating the probability at which an R-loop of a given (apparent) length arrives at a new length after time differences Δ*t* (as indicated). Shown are plots for data from measurements (upper row) as well as from Brownian dynamics simulations (lower row). Sketches on top illustrate intra- and intersegment R-loop length fluctuations at short and long time scales, respectively. **c,** Deconvolved free energy landscapes of R-loop formation for the different targets calculated from the apparent free energy landscapes in **a**. Shaded areas correspond to s.e.m. Dashed vertical lines indicate the flipped base positions. Numbers label the 5-bp duplex segments between the flipped base positions. **d,** Rate constants for the transitions between neighbouring 5-bp segments. **e,** Structural representation of the locked *E.coli* Cascade complex^27^ (PDB: 5he9) bound to the DNA target strand (left) together with a view on the crRNA – target strand hybrid (right). crRNA and DNA are shown in blue and green, respectively and the flipped DNA bases in purple. 5-bp segments are numbered as in **a** and **c**.

To gain insight into the dynamics of R-loop progression, we used transition plots that map the fate of R-loops of different lengths measured at a time *t* after certain delay times Δ*t* (Fig. 3b). For short delay times of 2 and 5 ms, the R-loop length became mainly redistributed within the energy wells, i.e. within a particular 5-bp segment, providing well localized maxima in the transition plots. For larger delay times, inter-segment transitions became allowed, i.e. the resulting R-loop lengths extended more and more to neighbouring wells. Thus, the R-loop length changes over longer time scales through 6-bp kinetic intermediate steps, as has been suggested from structural data^11^.

From the unwinding trajectories, we next determined the characteristic rate constants for intersegment transitions (Fig. 3d) between the 5-bp segments 2 to 5 (see Figs. 3a,e for numbering). For all target variants, we found similar intersegment transition rate constants ranging between 6 to 200 s^-1^. For a given pair of neighbouring segments, rate constants for R-loop extension were generally higher than for R-loop shortening due to the bias of the energy landscape. The rate constants for R-loop extension to segments 3 and 5 were particularly high, illustrating the more transient occupation of segments 2 and 4 compared to the more stable segments 3 and 5.

To obtain a realistic energy landscape for R-loop formation and quantify the free energy barriers between neighbouring segments, we set out to deconvolve the measured apparent energy landscapes. A direct deconvolution using the harmonic potential from the nanorotor fluctuations as a kernel provided unrealistically high energy barriers that would impede the R-loop dynamics (see Extended Data Fig. 9). To obtain more realistic energy landscapes, we used a maximum entropy deconvolution algorithm which reduced extreme energy barriers as function of an entropy scaling factor. Subsequently, the deconvolved energy landscapes were used in Brownian dynamics simulations that modelled the measurements of R-loop formation with the nanorotor assay (see Supplementary Discussions S6). From the simulated DNA untwisting trajectories, transition plots of the R-loop dynamics were constructed for a range of different entropy scaling factors as well as single base pair stepping rates of the R-loop. This allowed to identify the energy landscapes and the single base pair stepping rates that best described the measured R-loop dynamics (Fig. 3b, Supplementary Discussions S6).

The deconvolved energy landscapes exhibited more pronounced local energy wells and barriers with a 6-bp periodicity (Fig. 3c) compared to the apparent landscapes. Energy barriers between wells ranged from 0.7 *k_B_T* to 3.6 *k_B_T* for R-loop expansion and shortening, respectively (see Extended Data Fig. 10). Well minima were confined to only 1-2 bp, while energy barriers between wells were typically broader. This further supports a rather discrete segment-wise progression of the R-loop instead of a more continuous diffusion.

Mean-square-displacement analysis on short time scales showed good agreement between experiment and simulation (Extended Data Fig. 11) supporting that R-loop formation follows a single base pair stepping dynamics within the wells of the given energy landscape. We obtained single base pair stepping rates of 6000 ± 2000 s^−1^ (s.d.) corresponding to mean stepping times in the range of hundred microseconds. This is about 4-fold faster than stepping-rate estimates that neglected the periodic wells^16^ but about 10-fold slower compared to the stepping rates in strand displacement reactions of bare nucleic acid strands^15^.

We next investigated how a C:A mismatch between crRNA and DNA target strand at position 17 modulated the energy landscape (G17A mutation in the target strand). In theoretical models of DNA strand displacement reactions^15^ and target recognition by CRISPR-Cas effector complexes^14,16^, it has been previously hypothesized that a mismatch introduces a local free energy penalty for R-loop formation arising from the free energy penalty to generate the mismatch. Characterizing the R-loop formation dynamics on this substrate revealed two pronounced intermediates, one before and one after the mismatch where the latter one was less populated (Fig. 3a and Extended Data Fig. 8d). Reconstructing the free energy landscape for R-loop formation on the mismatch target revealed a pronounced free energy penalty for R-loop formation from positions 16 to 17 in agreement with the expectation. Our direct measurement of the energy landscape allowed an unambiguous determination of the penalty of 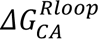 = 4.7 ± 0.5 *k_B_T* (Fig. 3c, Extended Data Fig. 8d and 10b), which was similar to the free energy cost of the same mismatch in a bare DNA duplex of 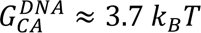. Notably, the presence of the mismatch affected only the R-loop transition rates between segments 3 and 4 (Fig. 3d and Extended Data Fig. 8d). This supports the idea that mismatches affect the R-loop dynamics only locally.

To explore the free energy landscape of R-loop formation beyond position 26 and probe its changes upon locking, we performed measurements on targets with fewer PAM-distal mismatches. While for the fully matching target the locking transition occurred too rapidly, targets with either two (T2) or one (T1) PAM distal mismatches allowed to sufficiently sample the free energy landscape of the unlocked R-loop at the PAM distal end (Fig. 4, Extended Data Table 2, Extended Data Fig. 12). Compared to the T6 target, the dominant well at 26 bp was broadened for the T2 and T1 targets and its minimum was shifted to an R-loop length of 27 bp. This means the R-loops were predominantly shorter than the fully matching regions of 30 and 31 bp for the T2 and T1 targets, respectively. Deconvolution correspondingly provided that the free energy landscapes of unlocked R-loops exhibited a pronounced uphill bias starting from position 27 despite of the absence of mismatches up to lengths of 30 and 31 bp, respectively (Fig. 4b).

**Fig. 4.**
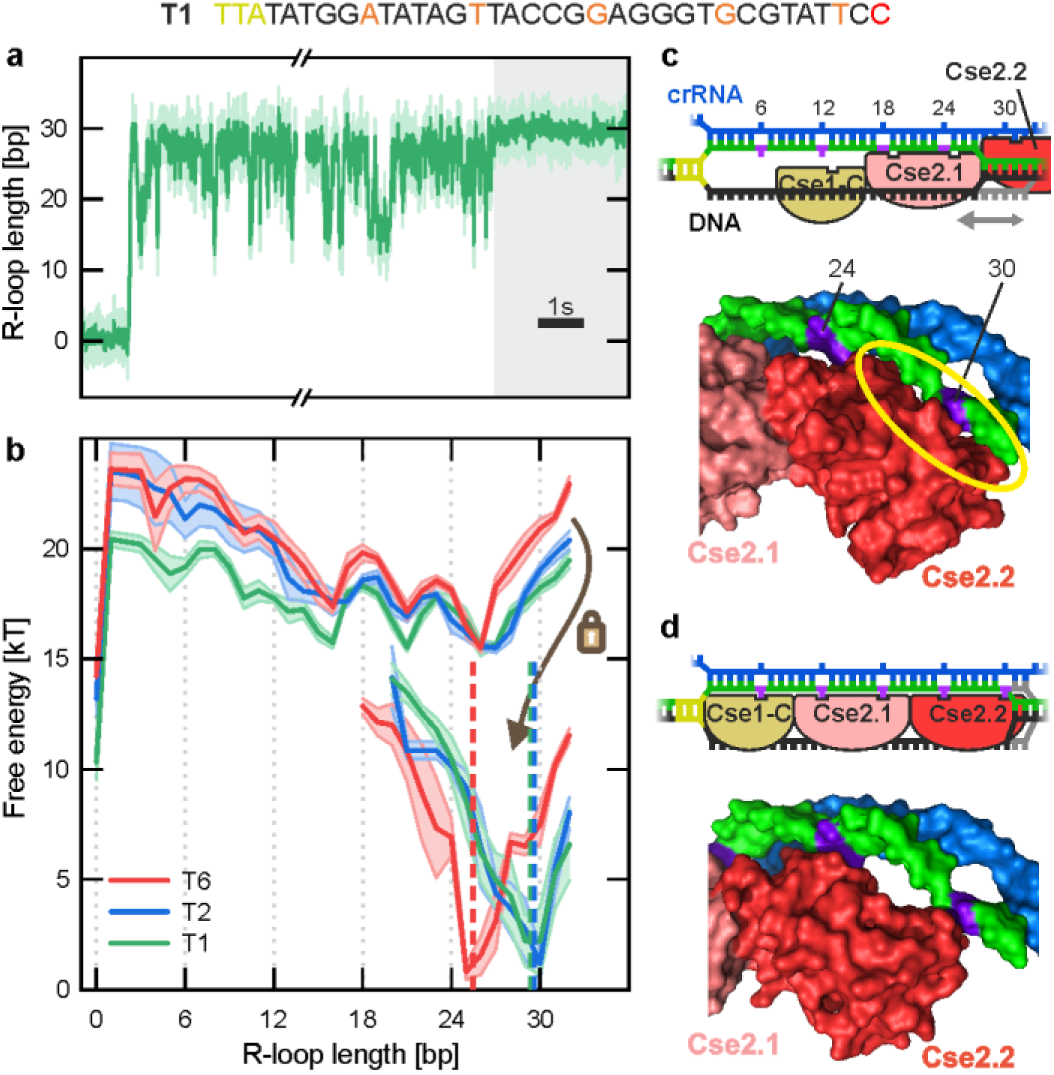
Locking requires R-loop formation against an uphill bias for lengths beyond 27 bp. **a,** Time trajectory of R-loop formation on a T1 target with a single PAM-distal mismatch. Following Cascade binding, highly dynamic R-loop length fluctuations were observed followed by sudden locking of the R-loop at a stable position (grey shaded area). The trajectory was recorded at 3947 fps (light colours) and smoothed with 100-point sliding average (dark colours) **b**, Deconvolved free energy landscapes of the unlocked (top) and the locked (bottom) R-loops measured for targets with six (T6), two (T2) and one (T1) PAM-distal mismatches. Shaded areas correspond to s.e.m. Dashed coloured lines indicate the dominant length in the locked state. The brown arrow indicates the transition into the locked state after reaching the full R-loop extension. Dashed grey lines indicate the flipped base positions. **c**, Structural representation of an overlay of the crRNA(blue) and Cse2 subunits (red) of the *E.coli* Cascade apo complex^28^ (PDB: 4tvx) with the target DNA (green) of the locked *E.coli* Cascade complex^27^ (PDB: 5h9e) revealing a steric clash between crRNA and Cse2.2 for R-loop length > 27 bp (yellow ellipse). The scheme above the structure illustrates that R-loop expansion beyond 27 bp is hindered by Cse2.2. **d**, Structural representation of crRNA, target DNA strand and Cse2 subunits of the locked *E.coli* Cascade complex^27^ (PDB: 5h9e). Sliding of the Cse2 subunits towards the PAM opens space for the hybridization of the target DNA and aligns its flipped bases with binding pockets in the Cse2 subunits as illustrated in the scheme above the structure.

The additional mismatches in case of the T6 target caused an earlier onset of the uphill bias. The uphill bias for R-loop lengths larger than 27 bp is in agreement with a steric clash of the DNA target strand with the Cse2.2 subunit of the unlocked Cascade complex starting at ∼28 nt from the PAM (see Fig. 4c), which was previously suggested based on the structures of *E.coli* Cascade complexes^29^. Since locked R-loop formation is highly sensitive to the number of PAM-distal mismatches^6^, we assume that the initiation of locking always requires the formation of a full R-loop of 32 bp length. The PAM-distal uphill bias together with mismatch penalties thus caused energy barriers of 7.4, 4.9 and 4.0 *k_B_T* that needed to be overcome for locking of the T6, T2 and T1 targets, respectively. Noticeably, the pronounced uphill bias provides similar free energy levels for the first and last base pair of the R-loop. While the initial negative bias would allow rapid R-loop formation on matching targets, the minimization of the total bias at the PAM-distal end would allow for a highly specific kinetic discrimination of mismatched targets^30^ (Extended data Fig. 12e,f).

After locking, the R-loop length became tightly constrained around 26 bp for the T6 and 30 bp for the T2 and T1 targets (Fig. 4, Extended Data Fig. 12). Upon locking, the filament formed by the Cse1 C-terminal domain, as well as Cse2.1 and Cse2.2 slides towards the PAM^27,29^. This provides space for full R-loop extension. Furthermore, the flipped bases of the DNA target strand get stably bound within pronounced pockets of the filament (Fig, 4d)^29^. Additionally, the R-loop gets further stabilized by the binding of the non-target strand to the backside of the Cse2 dimer^27^. In combination, those effects are thought to provide the high stability of the locked state. A highly confined R-loop of 30 bp for the T1 and T2 targets agrees with stable binding of the flipped base at position 30. The additional base pair that could form for T1 may not be stable in contrast to 2 bp for the fully matching target. The shallower slopes of the free energy landscapes towards lower lengths, suggest that locked R-loops of the T1 and T2 targets can still transiently shorten. The predominant R-loop length of 26 bp observed for the T6 construct can be understood by considering the additional mismatch penalties (Extended Data Fig. 12d). For this target the last flipped base would not be stably bound, albeit a local minimum in the free energy landscape of the locked state at 30 bp suggests at least transient R-loop extension and flipped base binding. Previous DNA twist measurements revealed that R-loop collapse proceeds via a transient intermediate in which 8-9 bp are rewound up to the flipped base at position 24^6^. This suggests that rupture of both flipped base contacts to the Cse2.2 subunit triggers R-loop collapse presumably since this provides the essential space for the backsliding of the Cse1-Cse2 filament into the unlocked state.

In summary, we constructed a nanomechanical rotor that enabled ultrafast twist as well as torque measurements on single DNA molecules over durations of hours. Key components of this assay were rigid DNA origami nanorotors combined with a high signal-to-noise detection of the rotor position using the large light scattering cross section of attached AuNPs. This allowed us to monitor in real time the R-loop dynamics within a Cascade effector complex and to reconstruct a detailed energy landscape for this process. R-loop formation is initially promoted by a weak global downhill bias up to a length of 27 bp but experiences a much steeper uphill bias for longer lengths. The landscape exhibits local energy minima distributed with a 6-bp periodicity, which directly reflects the structural periodicity of the crRNA-DNA hybrid. This forces the R-loop to expand with 6-bp kinetic intermediate steps despite of an underlying single base pair stepping on faster time scales. Full R-loop extension against the uphill bias triggers the repositioning of the Cse1-Cse2 filament due to a steric clash with the DNA target strand. Flipped base binding after locking, however, constrains the R-loop extension tightly and effectively quenches its dynamics.

We expect that our nanorotor assay will be directly applicable to other R-loop forming CRISPR-Cas effector complexes as well as to other proteins and enzymes that alter the DNA twist. Detailed insight into the R-loop formation dynamics and energetics will be important to work out the different mechanisms by which CRISPR-Cas effector complexes achieve their specificities^30^ and to establish and validate mechanism-based off-target predictions for these enzymes^14,16,31^.

## ACKNOWLEDGEMENTS

This work was supported by grants from the European Research Council (GA 724863) and the Deutsche Forschungsgemeinschaft (SPP2141, SE 1646/8-1) to R.S.. J.M.M was supported by a postdoctoral fellowship of the Alexander von Humboldt Foundation. We thank Jingjing Ye and Ulrich Kemper for providing TEM images of the nanorotor as well as David Poppitz for the training and support of the TEM imaging. We further thank Patrick Irmisch and Fabian Welzel for critical reading of the manuscript.

## AUTHOR CONTRIBUTIONS

R.S. and D.J.K. designed the study. I.S., T.S. and V.S. provided the purified Cascade. D.J.K. and A.W. designed the nanorotor. M.R. designed the DNA targets. D.J.K., J.M.M. and A.W. carried out the experiments. D.J.K. analysed the data. D.J.K., J.M.M. and M.R. interpreted the results. D.J.K and R.S. and wrote the manuscript. All authors provided comments to the manuscript.

---

**Extended Data Table 1.** Nanomechanical parameters and spatio-temporal resolution of different single molecule rotor assays.

| Value | AuNP nanorotor<br>this study | AuNP rotor bead<br>applied to enzymes<br>(Ivanov et al.<sup>22</sup>) | Fluorescent nanorotor<br>(Kosuri et al.<sup>21</sup>) |
| --- | --- | --- | --- |
| $\kappa$ [pN nm / rad] | $5.0 \pm 0.2$ | $\approx 1.7$ | 7.8 |
| $\gamma$ [fN nm s] | $6.5 \pm 0.4$ | $\approx 5$ | 5.0 |
| RMS [rad] | $0.91 \pm 0.02$ | $\approx 1.6$ | 0.72 |
| $\tau$ [ms] | $1.3 \pm 0.1$ | $\approx 2.9$ | 0.64 |
| $T_{1\text{ bp}}$ [ms] | $54 \pm 4$ | 410 | 17 |

Shown are the torsional trap stiffness $\kappa$ , drag coefficient $\gamma$ , root mean square (RMS) amplitude, response time $\tau$ and averaging time $T_{1\text{ bp}}$ required to reach a resolution of 1 bp with SNR = 3. Values for our study represent averages from 18 molecules given as mean ± s.e.m. A more detailed comparison of the different rotor assays is given in Supplementary Discussions S3.

**Extended Data Table 2.**
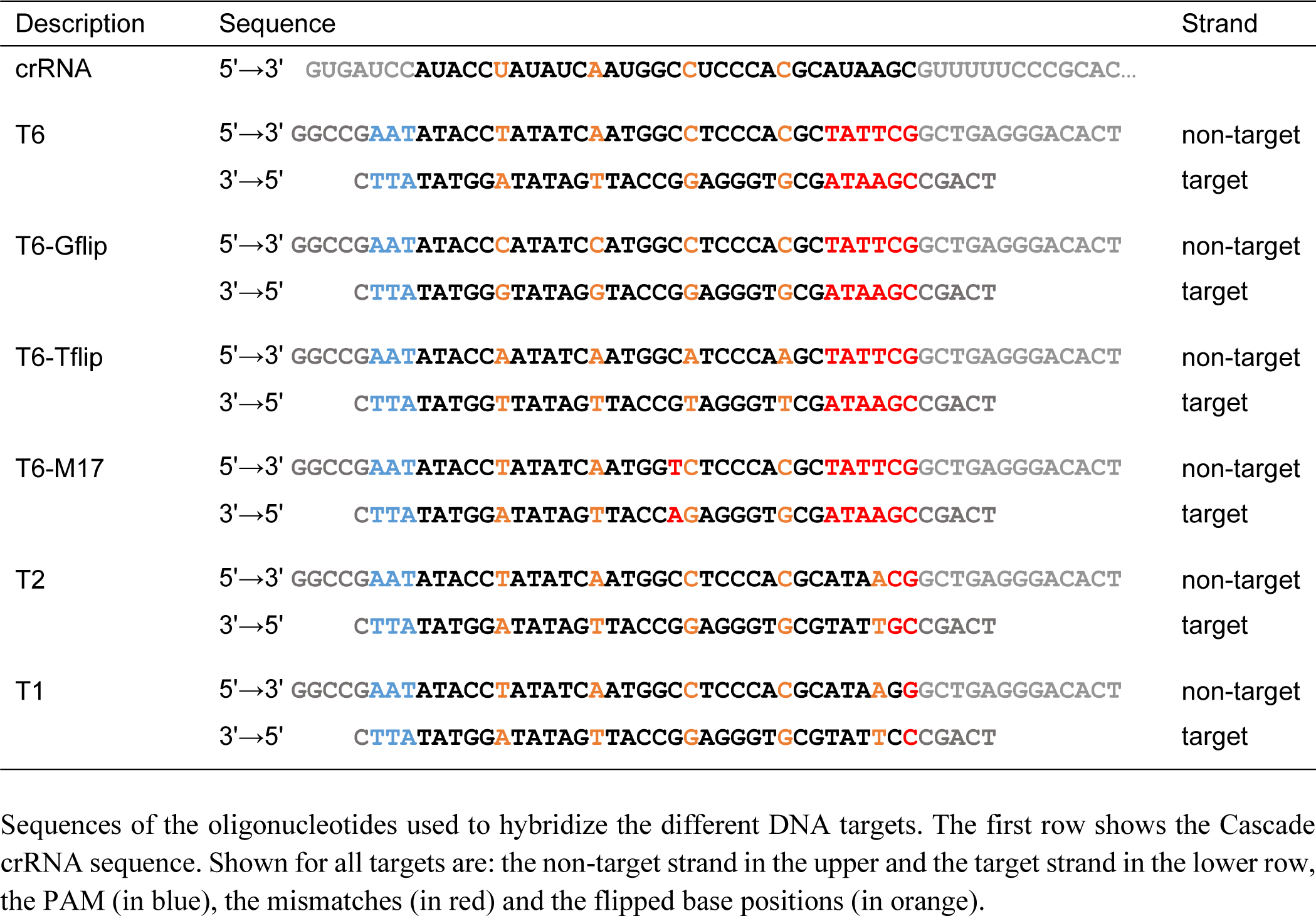
Target sequence oligonucleotides.

**Extended Data Table 3.**
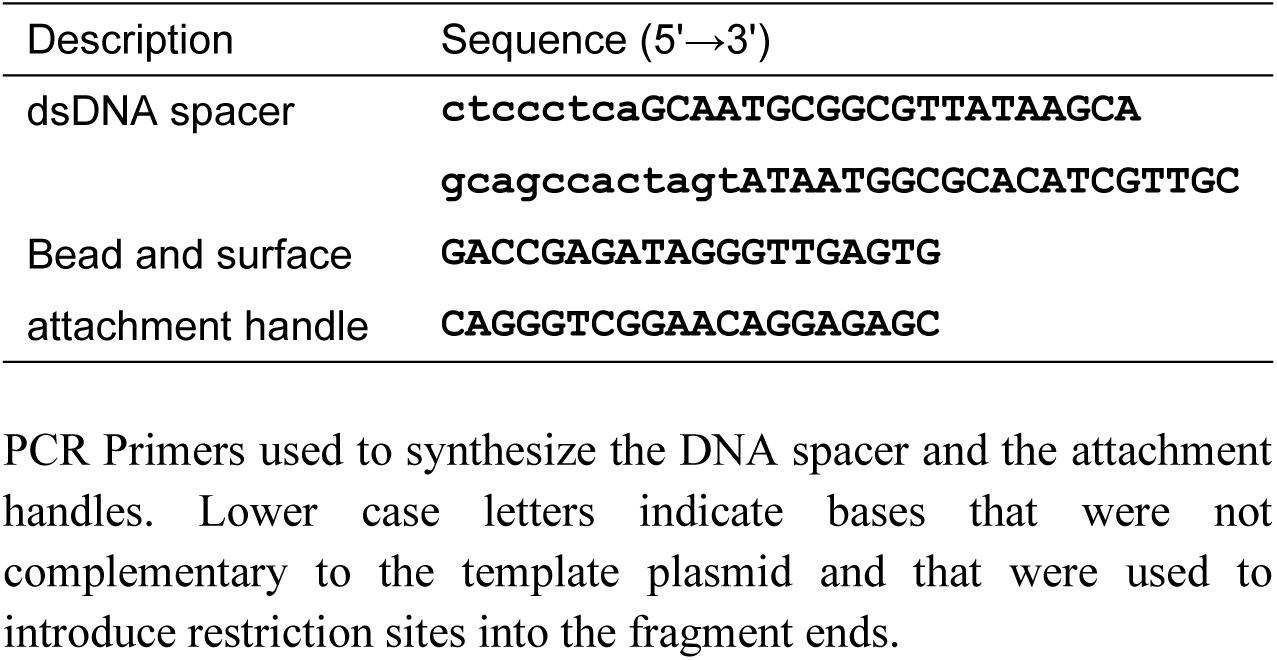
PCR primer sequences.

**Extended Data Table 4.** DNA quantities used in the ligation of the nanorotor construct.

| Element | Amount [fmol] |
| --- | --- |
| Bead attachment handle | 400 |
| dsDNA spacer | 50 |
| Nanorotor | 75 |
| Target sequence | 200 |
| Surface attachment handle | 800 |

**Extended Data Fig. 1.**
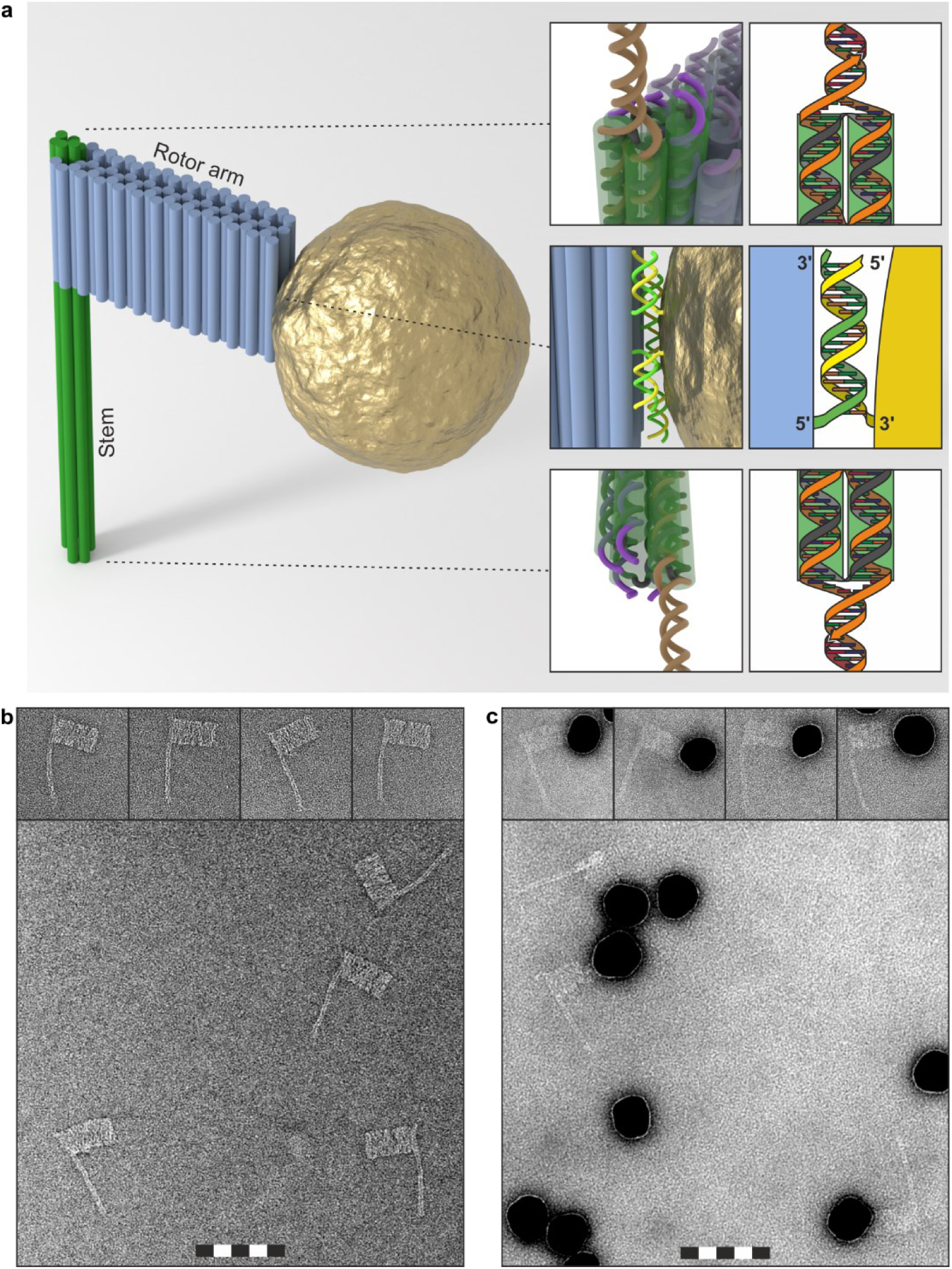
Scheme and TEM images of the nanorotor. **a,** Three-dimensional scheme of the nanorotor including the origami rotor arm and a 50 nm AuNP. Blue cylinders represent the individual dsDNA helices. Insets at the top and bottom show enlarged perspectives as well as schematic views of the dsDNA ligation interfaces at each end of the stem, where two complementary ssDNA staple overhangs of neighbouring helices (orange) and a secondary interface oligo form a sticky end for ligation. Other helix ends carry six nucleotide ssDNA staple overhangs (purple) to prevent aggregation between rotor arm structures due to blunt end DNA stacking. The insets in the middle show detailed and schematic views of the attachment of the AuNP to the end of the rotor arm, in which the connecting DNA strands hybridize in a zipper-like configuration. **b,** TEM overview image of several DNA origami rotor arms and selected images of individual structures (top). **c,** TEM overview image of DNA origami rotor arms with attached 50 nm AuNPs and selected images of individual structures (top). The nanorotor binds often to protrusions on the AuNP. All TEM images are scaled to the same magnification. The length of the scale bars is 100 nm with 20 nm for the length of each black and white segment.

**Extended Data Fig. 2.**
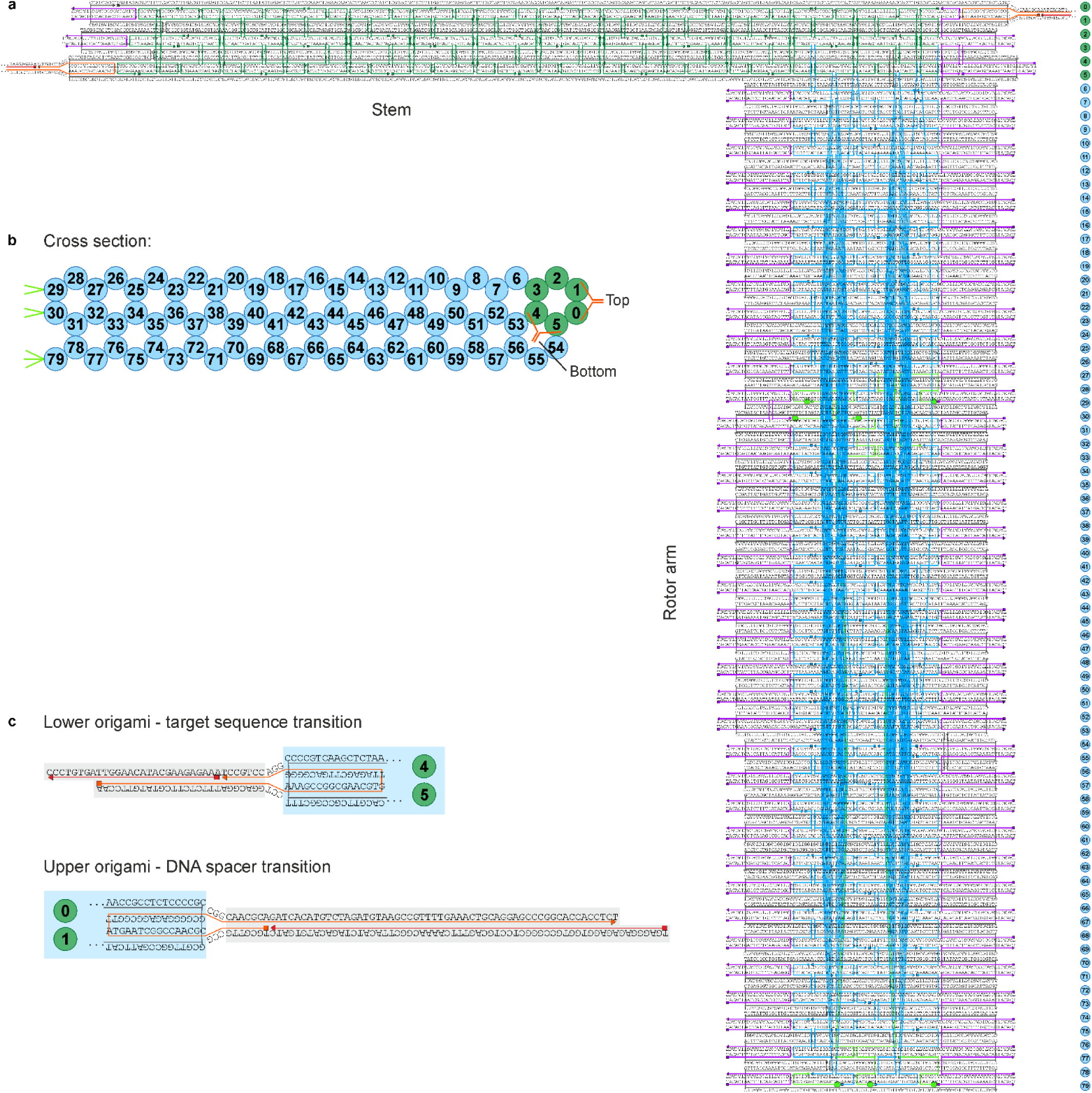
Design scheme of the DNA origami rotor arm structure. **a,** An 8064 nt ssDNA scaffold (black) is assembled into the DNA nanostructure in a reaction with 206 staple oligonucleotides. The stem is made of a 6-helix bundle (green staples), from which a rotor arm emerges (blue staples). Staples at duplex ends have six nucleotide ssDNA (“TACACT”) overhangs (purple) to prevent aggregation of the origami structures via blunt end stacking interactions^32^. On each end of the stem, two ssDNA overhangs of two neighbouring helices (orange staples) can hybridize to form a terminal duplex with a sticky end for ligation to the adjacent dsDNA parts. Seven staple oligonucleotides that terminate with their 3’-end at the stem-distal end of the rotor arm (shown in bright green, helices 29, 30 and 79) carry 20 nt poly-A 3’ overhangs (bright green circles) to allow the attachment of an AuNP coated with 20 nt 5’-SH-poly-T oligonucleotides (see Extended Data Fig. 1). The total length of the stem is 106 nm; the total extension of the lever arm from the stem is 56 nm, assuming an inter-helical distance of 2.536 Å^33^. b, Top view on the rotor arm design showing the arrangement of the parallel DNA helices on a honeycomb lattice. The helices forming the stem are shown in green, other helices in are shown in blue. Overhangs for AuNP attachment are shown as green lines. Y shaped orange markers show the exit points of the connector oligonucleotides from the stem that allow the connection of the adjacent dsDNA parts. c, Detailed design of connector oligonucleotides at the stem ends. The primary interface oligo (orange) is incorporated during the assembly of the DNA origami structure. It can weakly hybridize to itself over a few base pairs. In a second step, a secondary interface oligo (red) is annealed to the overhang of the primary oligonucleotide such that an 8 nt sticky end for the subsequent ligation is formed. After ligation, the connector oligonucleotides form a loop which is topologically locked with the DNA scaffold at the stem ends to support a high twist load.

**Extended Data Fig. 3.**
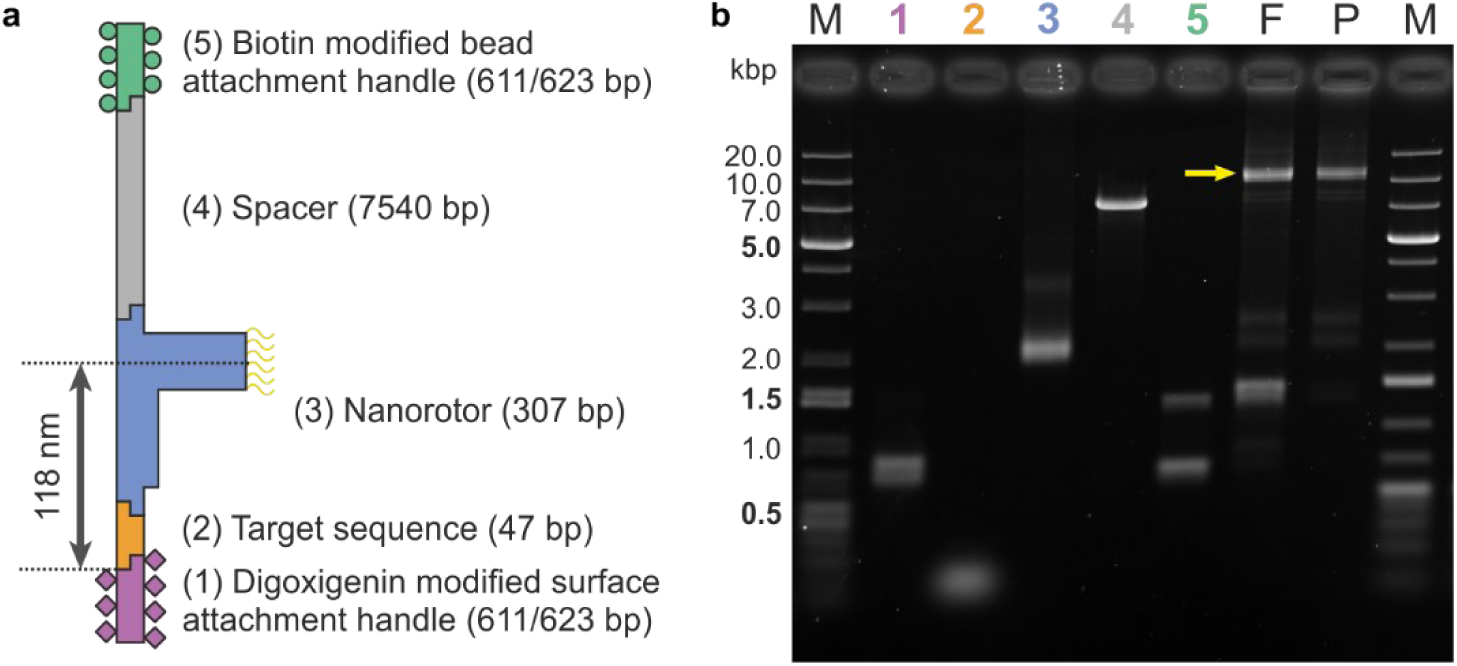
Design and construction of the nanorotor DNA construct. **a,** Scheme of the construct components including four dsDNA fragments: a digoxigenin-labelled surface attachment handle (1), the Cascade target sequence (2), a long spacer to lift the magnetic bead out of the evanescent field (4) and a biotin-modified bead attachment handle (5), as well as the origami rotor arm structure that carries attachment sites (yellow) for the gold nanoparticle (3). **b,** Analysis by agarose gel electrophoresis of the individual components of the DNA nanorotor construct (lanes 1-5, numbered according to a) as well as the ligation products from all components before (lane F) and after PEG purification (lane P). Lane M contains a DNA size marker whose dsDNA fragment length is given on the left. In lane F, a yellow arrow marks the band containing the full nanorotor DNA construct. Additional dominant ligation products at 1.0-1.5 kbp are caused by the self-ligation of both handles due to the use of an excess of handles with symmetric ligation overhangs. Low molecular weight products (< 1.5 kbp) were removed in the subsequent PEG precipitation step. Remaining side-products were removed upon magnetic bead binding and washing prior to the tweezers experiments. Gel electrophoresis was performed as described previously^34^.

**Extended Data Fig. 4.**
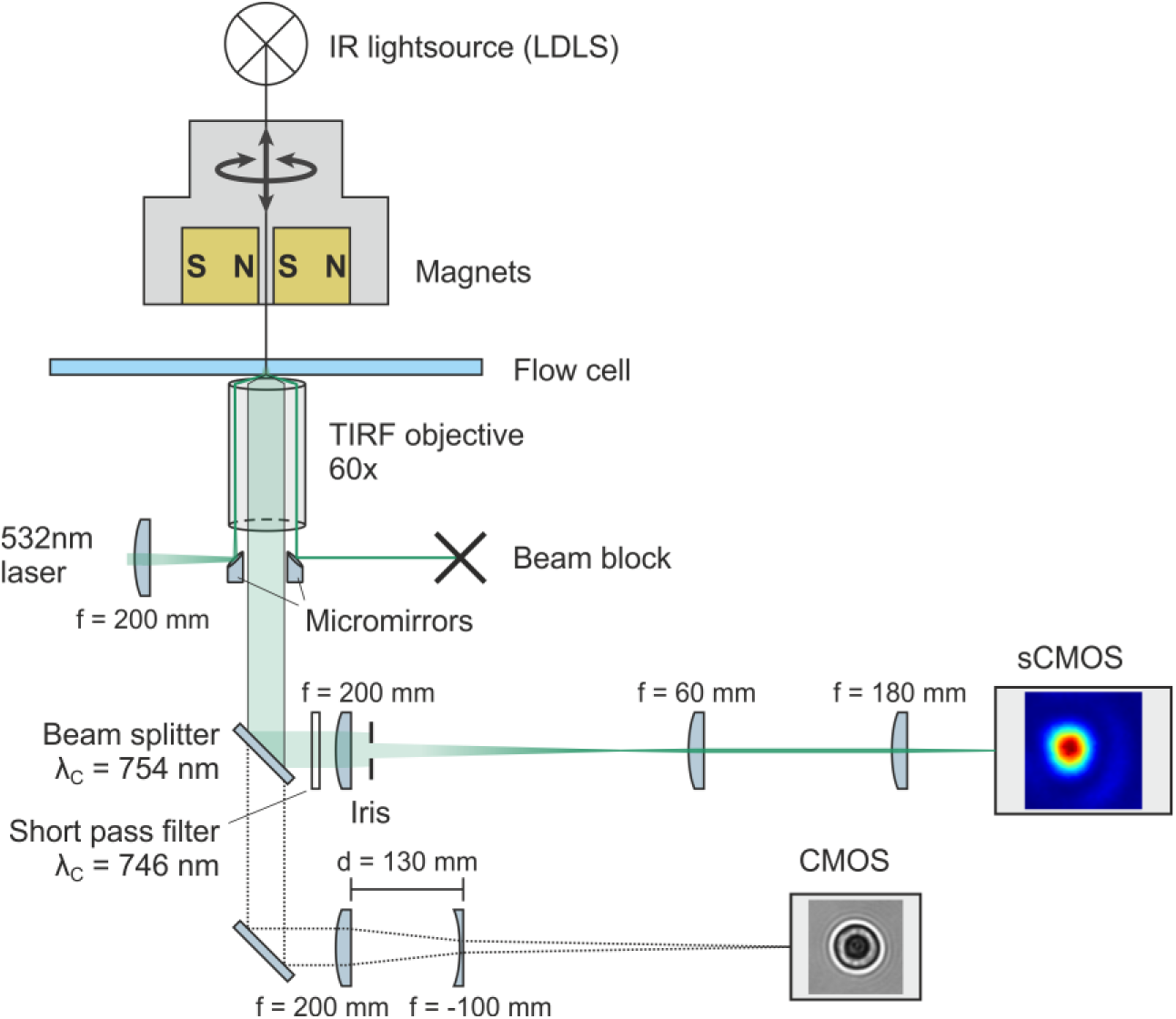
Setup combining magnetic tweezers with light scattering microscopy. A green laser (λ=532 nm) is focused via an external lens and a micro mirror into the back focal plane of a high-numerical aperture oil-immersion objective (CFI Apochromat TIRF 60XC Oil, Nikon, Japan). Laterally displacing the incident beam from the optical axis allows to illuminate the sample in total internal reflection geometry. Specifically, an evanescent field is generated that exponentially decays from the surface of the bottom cover slip into the aqueous medium with a depth of ∼200 nm. The reflected laser light is coupled out of the microscope using another micro mirror. Light that is scattered inside the flow cell (e.g. at the AuNP of the nanorotors) passes the objective pupil between the micro mirrors and is projected onto a sCMOS camera (C11440-22CU, Hamamatsu Photonics). Images of the scattered light were recorded at 3947 fps. For magnetic bead tracking, the flow cell is illuminated by IR light using the focused beam of a strong, non-coherent laser-driven arc lamp (LDLS, EQ-99, Energetiq Technology, Wilmington, MA) which was spectrally filtered (770−950 nm wavelength). The IR light from the LDLS is separated from the green scattered light using a beam splitter (cut-off at 754 nm wavelength). Using a suitable tube lens, it is then projected onto a CMOS camera (EoSens CL MC-1362, Mikrotron, Germany), which records images of selected magnetic beads at 1000 fps. A pair of permanent magnets above the flow cell is used to generate a strong magnetic field gradient inside the flow cell. The magnets can be rotated around the optical axis and moved vertically, which allows to apply twist and stretching forces onto the magnetic beads and the attached DNA nanorotor constructs (Fig. 1, Extended Data Fig. 7).

**Extended Data Fig. 5.**
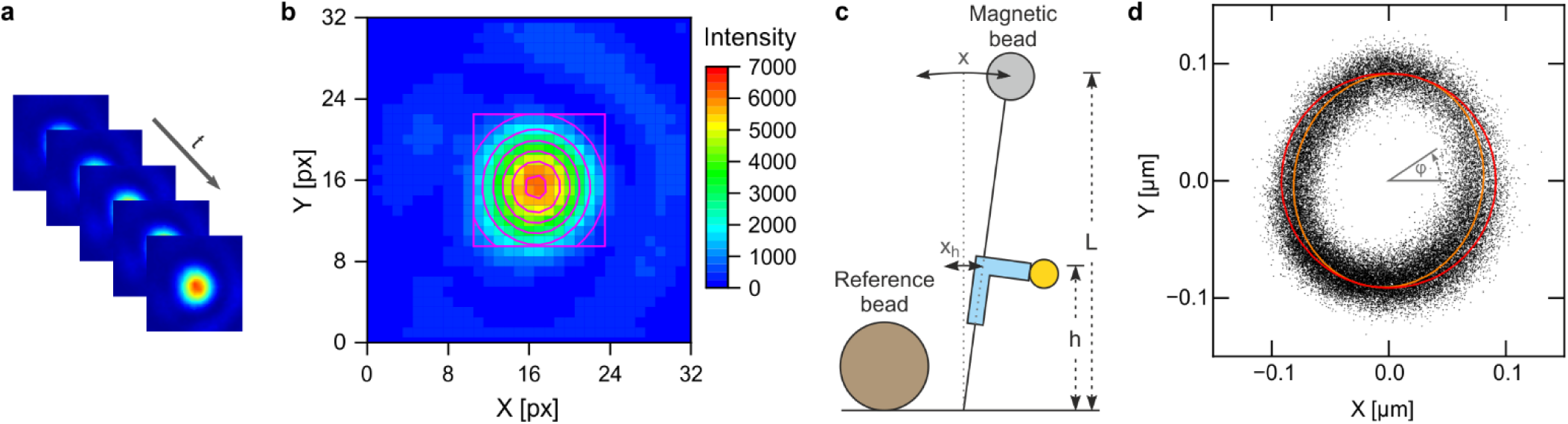
Tracking the motion of nanorotor AuNPs and determination of the angular position. **a**, Stack of consecutive images of an individual 50 nm AuNP attached to a DNA nanorotor construct recorded at 3947 fps. The image size is 32 px x 32 px corresponding to a field of view of 1.32 µm. **b,** Localization of the AuNP. To determine the position of the AuNP centre, the intensity distribution of a 13 px x 13 px area (pink square) around the pixel with maximum intensity was fitted with a two-dimensional Gaussian function (pink circles, indicating Gaussian intensities of 6000, 5000, 4000, 3000, 2000, 1000 from inside to outside). Measurements on AuNPs stuck to the glass substrate yielded a resolution of 2.7 nm. **c,** Corrections for magnetic bead fluctuations and drift of the microscope. In parallel to the tracking of the AuNP, we record the positions of the attached magnetic bead and a surface-attached non-magnetic reference bead. The position tracking of the reference bead is used to remove the thermal drift of the microscope. It is applied within a real-time z-drift correction and a coarse lateral-drift correction that prevents the AuNP from leaving the small field of view. During off-line tracking of the AuNP position, the reference bead position is used for a precise lateral-drift correction. Thermal fluctuations cause lateral displacements *x* of the magnetic bead from the equilibrium position. In turn this results in lateral displacements *x_b_* of the nanorotor rotation centre. To correct for this, we determined *x_b_* from *x_b_* = ℎ⁄*L* ⋅ *x*, where *L* is the distance of the magnetic bead centre from the DNA surface attachment (obtained from bead tracking) and ℎ is the height of the AuNP above the surface for which 118 nm was taken. **d,** Tracked positions of a nanorotor bound AuNP in the XY plane. Imperfections of the magnet configuration can cause a tilt of the pulling direction with respect to the optical axis, providing an elliptic distortion of the AuNP positions due to the tilt of the rotation plane. A fit of the data with an ellipse is shown in orange. To correct for the tilt of the rotational plane, the short ellipse axis was stretched to the value of the long axis to yield a circular trajectory (shown in red). This correction was applied to all data points in order to obtain correct angular positions of the nanorotor.

**Extended Data Fig. 6.**
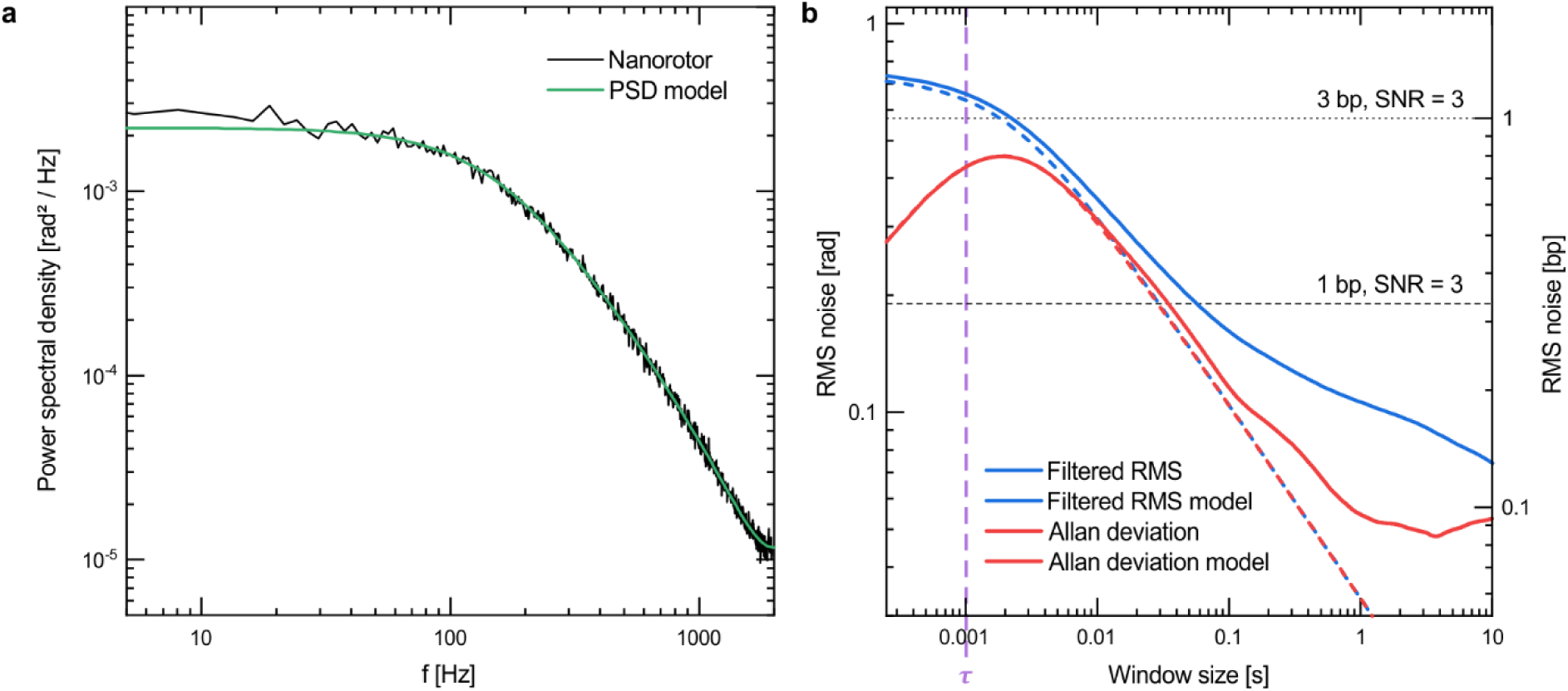
Power Spectral Density and Allan Deviation of nanorotors. **a,** Power spectral density (PSD) of a nanorotor (black) that was torsionally constrained on the bottom side (next to the Cascade target) in absence of Cascade. The plateau seen at low frequencies in absence of Cascade demonstrated that the angular nanorotor fluctuations were considerably confined. This confinement was released when dynamic R-loops formed in presence of Cascade (see Fig. 1d, Fig. 2a). The PSD of the Cascade-free trajectory was fitted with a Lorentzian that was further corrected for aliasing and signal integration function (green, see Supplementary Discussions S1). Best-fit parameters for the shown molecule were *k_rot_* = 7.5 ± 0.1 pN nm/rad, *f_cut_* = 158 ± 1 Hz). **b,** Allan deviation (red solid line), calculated using overlapping intervals^35^ and root mean square (RMS) noise after filtering with a sliding average (RMS, blue solid line) of the nanorotor trajectory used in **a** (in rad and in bp). Dashed lines of the same colour represent theory predictions for the best-fit parameters of the PSD (see Supplementary Discussions S2). The response time of the nanorotor was τ = γ/*κ* = 1.0 ms (purple dashed line). Horizontal dashed lines indicate the noise level at which DNA twist changes of 3 and 1 bp become resolvable assuming a SNR of 3. Intersections of this lines with measured RMS noise provide the corresponding temporal resolution to detect these twist changes of ∼4.5 ms for 3 bp and ∼54 ms for 1 bp.

**Extended Data Fig. 7.**
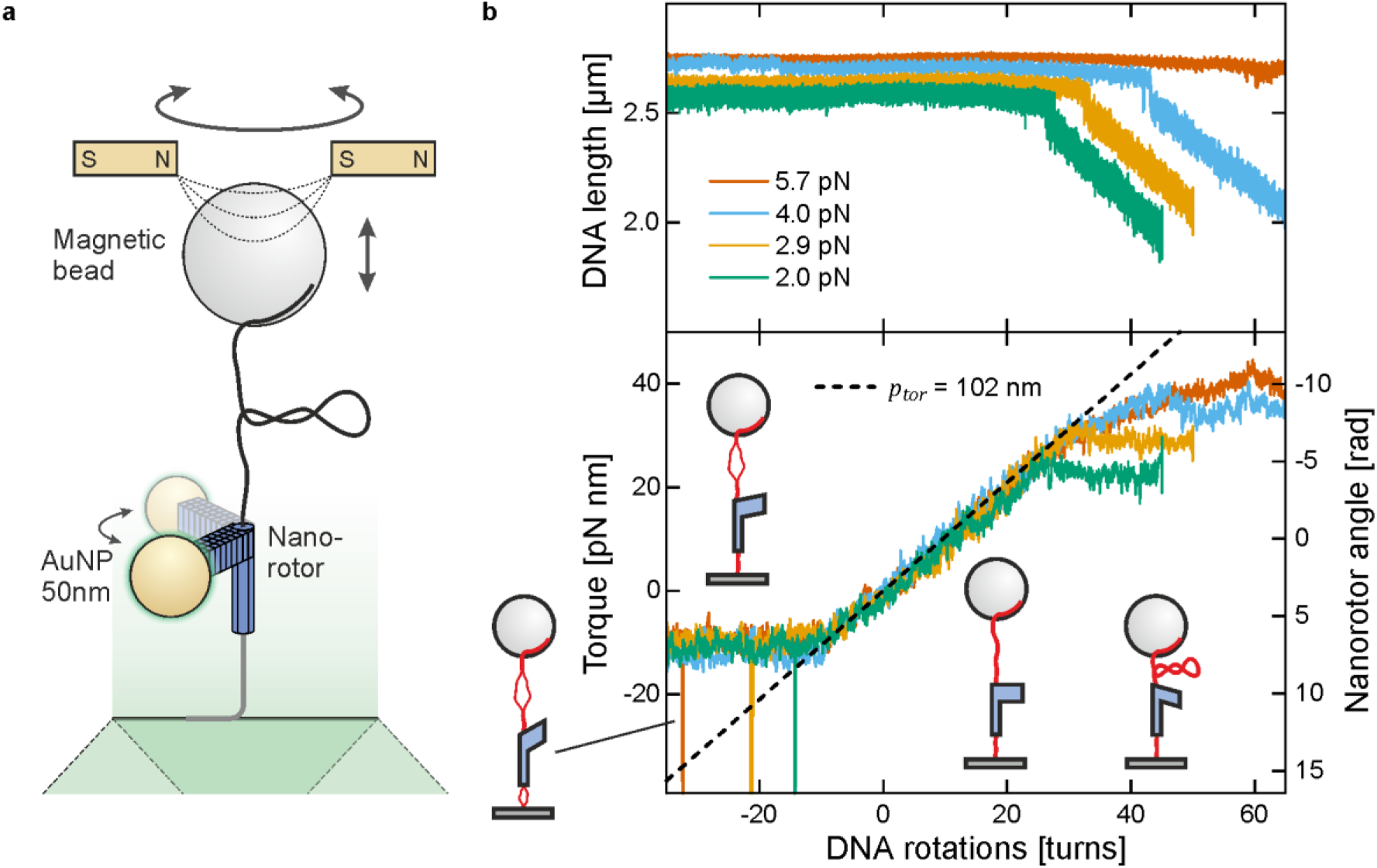
Direct torque measurements during DNA twisting using the nanorotor. **a,** Experimental setup for torque measurements during DNA twisting. Torque measurements require the nanorotor to be torsionally constrained with respect to both attachments, i.e. at the surface and at the bead. Twisting the DNA by magnet rotations changes the torque in the molecule, which in turn displaces the nanorotor. Calibrating the torsional stiffness *k_tot_* of the nanorotor system (see Extended Data Fig. 6), which is dominated by the linker on the bottom, allows to obtain the torque from the angular displacement of the nanorotor (see Supplementary Discussions S4). **b,** DNA length and torque during DNA twisting at different forces. Around zero turns, the DNA length remains approximately constant, while the torque is increasing proportionally with the applied turns. From the slope of this linear part, a torsional persistence length of dsDNA of *p_tor_* = 102 nm was obtained, which is in good agreement with previous reports^36^. Once a critical positive torque was reached, the DNA buckles, which is seen as a sudden length jump. This is followed by a linear DNA length decrease with applied turns in which the plectonemic writhe structure (see sketch in torque plot) is further expanding. In the plectonemic regime, the torque remains approximately constant. Both the buckling point as well as the torque plateau are force dependent. For the measurement at 5.7 pN, no length decrease is observed, as the DNA undergoes a transition to p-DNA^25^. When the negative torque falls below −9 pN nm, another torque plateau is reached, where additionally applied negative turns are absorbed by force-independent local melting or Z-DNA formation^37^. These local structural transitions appear to be dynamically changing throughout the molecule. They are typically found in the long DNA spacer above the nanorotor but can transiently appear as well in the dsDNA part below the nanorotor (see transient spikes towards negative nanorotor angles). Measurements are performed at a resolution of 1 pN nm within 38 ms which allowed to measure the torque continuously while twisting, instead of averaging measurements at set twist values^38^.

**Extended Data Fig. 8.**
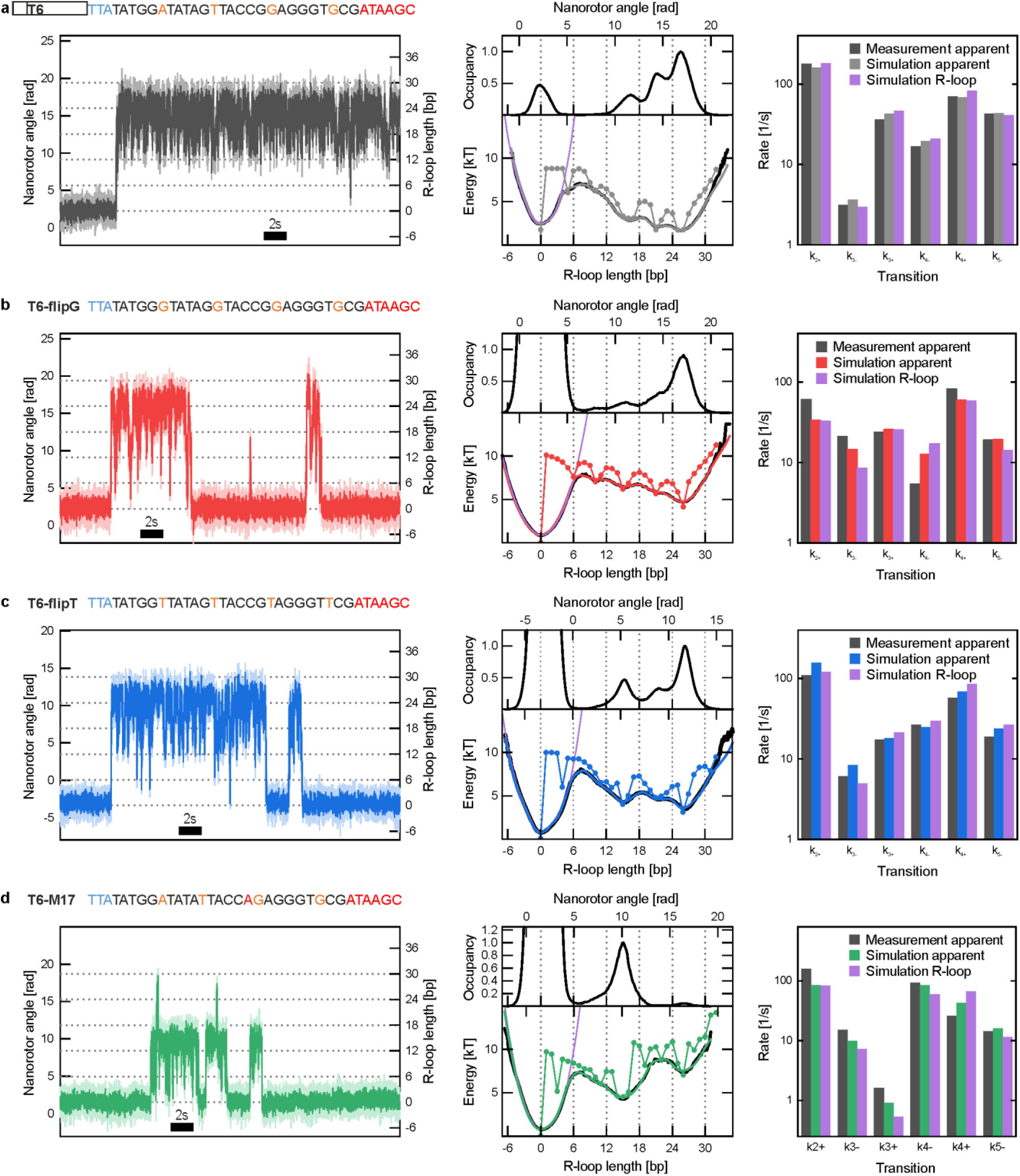
Trajectories, energy landscapes and inter-well transition rates for R-loop formation on the different T6 target sequences. Shown are results measured on a single molecule with a, T6, b, T6-flipG, c, T6-flipT and d, T6-M17 target sequence. For each target are shown: a DNA untwisting trajectory in presence of Cascade (left, light colours represent raw data at 3947 fps, dark colours the data after 100-point adjacent averaging), the sequence of the particular target (with PAM bases in blue, the flipped base positions in orange and mismatches with the crRNA in red), a histogram of the angular positions obtained from all trajectories of the given molecule (middle, top) as well as the corresponding free energy landscapes for R-loop formation (middle, bottom). In the latter, the apparent (black line), the deconvolved (line and scatter points) and the reconvolved (coloured line) free energy landscapes are shown. The apparent energy landscape in absence of Cascade (from which the deconvolution Kernel was obtained) is shown as a purple line. Inter-well transition rates from measured and simulated DNA untwisting trajectories are shown on the right. In the simulations, the R-loop length can be monitored directly from which correct transition rates of the simulations were calculated. Transition rates obtained by the three different ways show a good overall agreement. This validates the determination of realistic inter-well transition rate from experimental trajectories.

**Extended Data Fig. 9.**
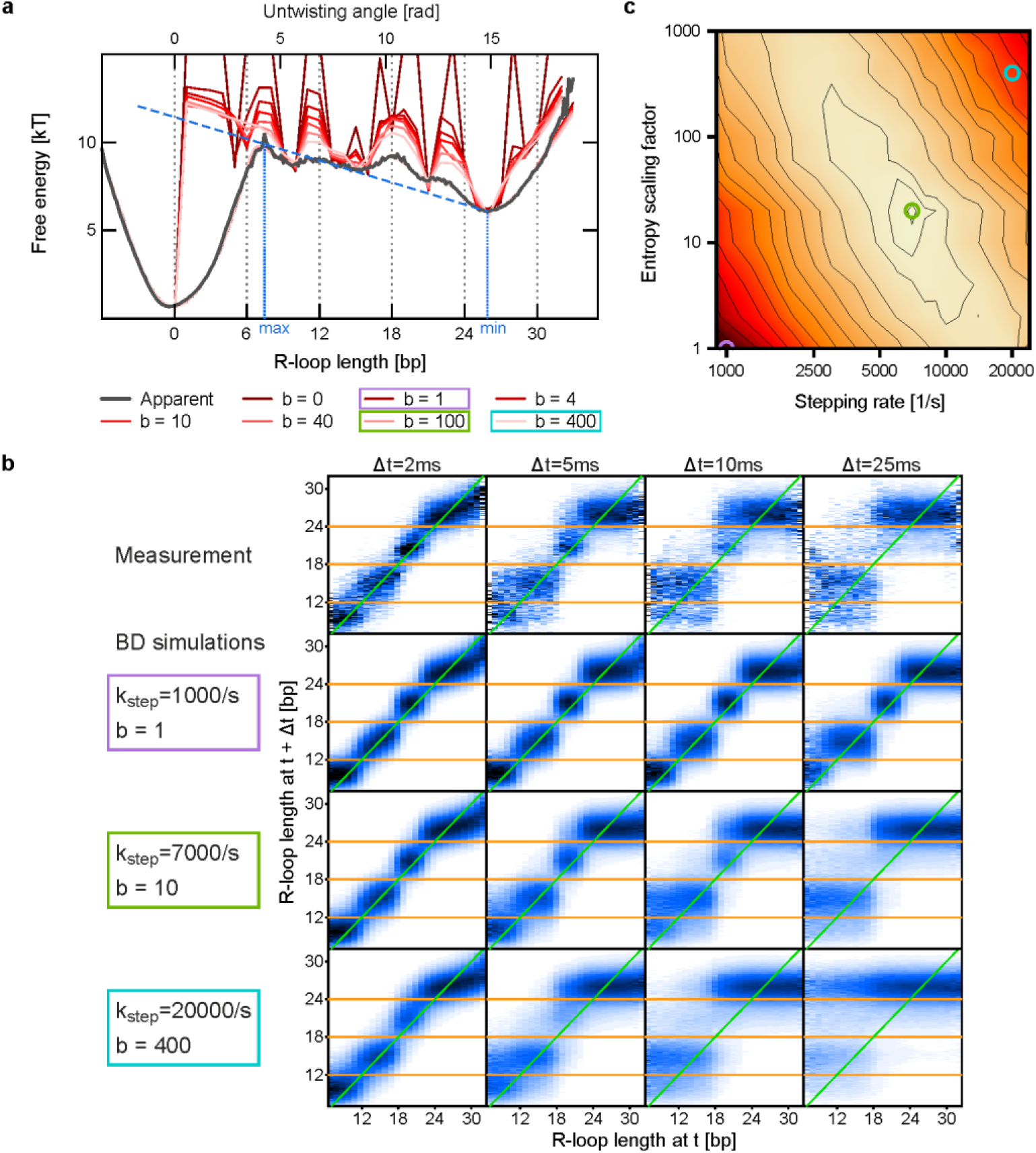
Deconvolution of an apparent free energy landscape of R-loop formation exploiting the measured R-loop dynamics. **a,** Apparent (black line) and deconvolved (red lines) free energy landscape of R-loop formation obtained for the T6-flipG target. Deconvolved energy landscapes were calculated for different entropy scaling factors *b* (see legend). The blue dashed line is a linear approximation of the apparent energy landscape between the local maximum at ∼7 bp and the minimum at 26 bp. Increasing entropy scaling factors decrease the height of the free energy barriers between the local energy wells. To obtain a deconvolved free energy landscape that supports the measured dynamics of the R-loop formation as well, transition probability plots were used (see b). b, Transition maps revealing the dynamics of R-loop formation. Each transition map shows the probability distribution describing the length change of R-loops (vertical axis) of a given apparent length at time *t* (horizontal axis) after various time delays Δ*t* (2, 5, 10 and 25 ms). Shown are maps for the measurement in a (upper row) as well as for the Brownian dynamics simulations using the deconvolved energy landscapes obtained for different energy scaling values *b* and single base pair stepping rates *k_step_*. While all *b* values well describe the intra-segment diffusion at 2 ms, significant differences are seen for the inter-segment dynamics at larger time scales, for which *b* = 10 and *k_step_* = 7000 *s*^−1^ provide the parameter fit that best describes the measured data. c, Total residue (RMS) between simulated and measured transition plots (for all time differences) shown for different parameter sets of *b* and *k_step_*. The RMS for a given parameter set was calculated from the sum of the RMS values at Δ*t* = 2, 5, 10 and 25 ms). Coloured circles correspond to the 3 parameter sets highlighted/chosen in a and c. Across all targets we obtained a single base pair stepping rate of *k_step_* = 6000 ± 2000 *s*^−1^ (s.d.).

**Extended Data Fig. 10.**
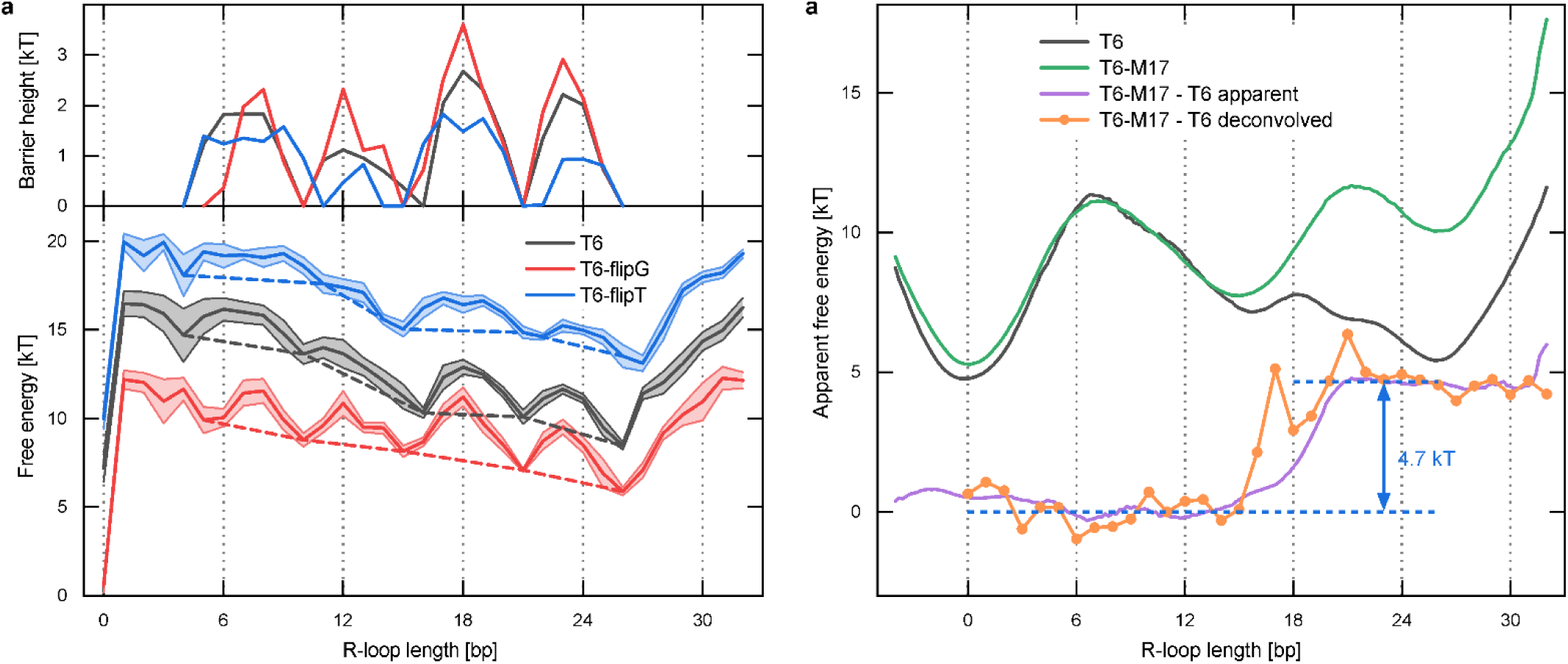
Height of intersegment and mismatch barriers in the energy landscape. **a,** (Top) Free energy barriers for intersegment transitions for the T6, T6-flipG and T6-flipT targets (see legend). (Bottom) The height of the transition barriers of a particular free energy landscape was determined from its difference to a barrier-free reference obtained by connecting the minima of adjacent free energy wells (dashed lines). The resulting barriers were approximately centred on the flipped base positions (dotted grey lines). **b,** Free energy penalty introduced by a single C:A mismatch at position 17 between the target strand and crRNA. Shown are the apparent energy landscapes of the T6 (black) and the T6-M17 (green) targets as well as the difference between both apparent (purple) and both deconvolved (orange) energy landscapes. The difference between the two landscapes reveals a pronounced mismatch penalty, i.e. an offset of the energy values for lengths larger 17 bp, of 4.7 ± 0.4 *k_B_T* (see difference between the blue dashed lines).

**Extended Data Fig. 11.**
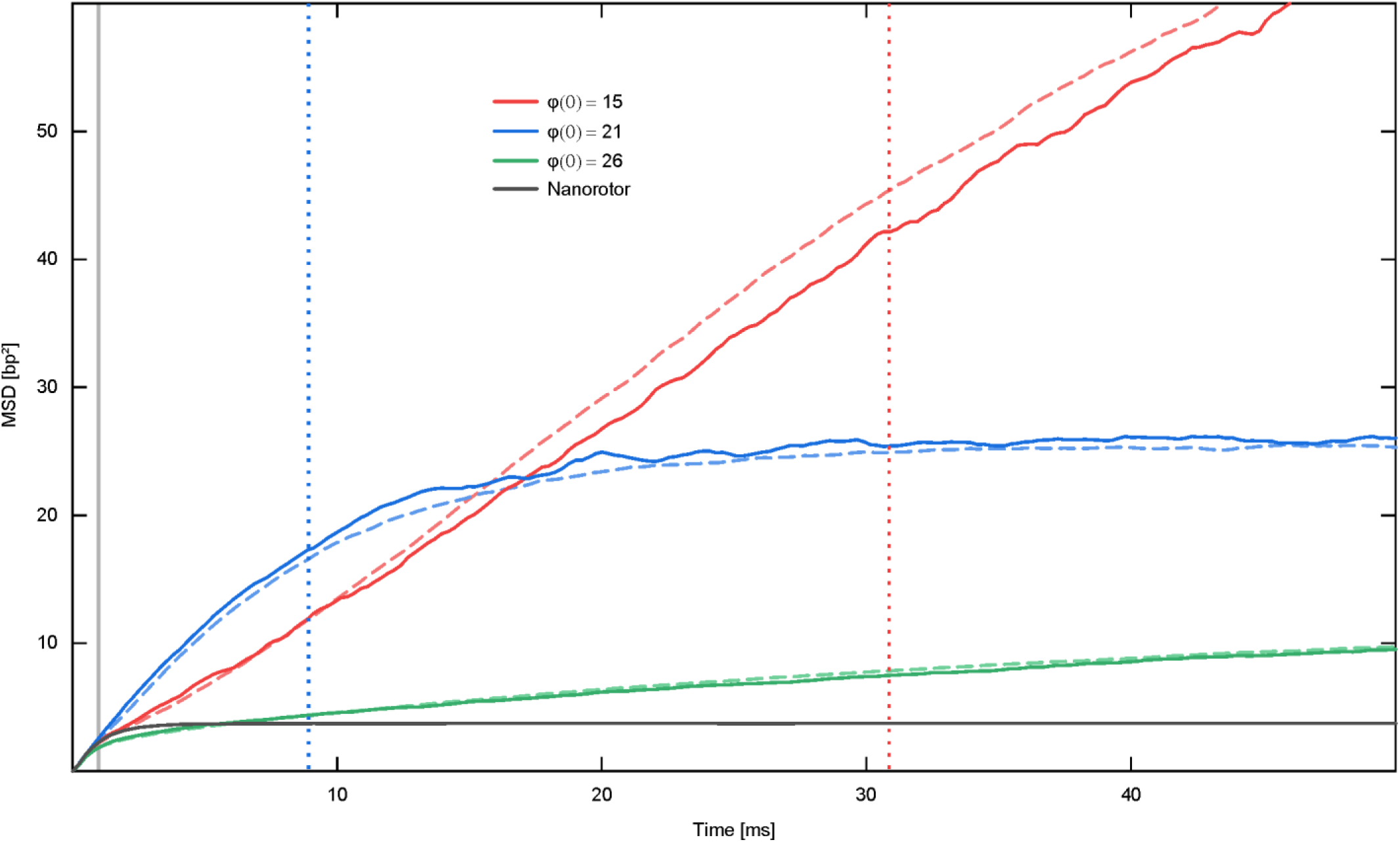
Mean-squared-displacement (MSD) of R-loop length fluctuations over time in comparison to simulations. **a,** Mean squared displacement of the bare nanorotor (black) as well as of the R-loop length fluctuations as a function of time 〈(φ(*t*) − φ(0))^2^〉 for starting positions φ(0) of 15, 21 and 26 bp corresponding to the 5-bp segments 3, 4 and 5, respectively. Shown are MSDs from the measurement on a molecule with a T6 target (solid lines) and Brownian dynamics simulation using the best-fit parameters: *b* = 20 and *k_step_* = 7000, for its R-loop dynamics (dashed lines). The measured mean escape times (dotted vertical lines) from the wells 3, 4 and 5 were 31, 8.9 and 74 ms respectively. The good agreement between measurement and simulation for the MSD plots provides strong support for the obtained energy landscape and the stepping rate *k_step_*. In particular, the agreement for short times in between the nanorotor response time (grey vertical line) and the corresponding escape times from the wells strongly supports that the intrawell R-loop dynamics is governed by a random walk-like process of 1 bp step size.

**Extended Data Fig. 12.**
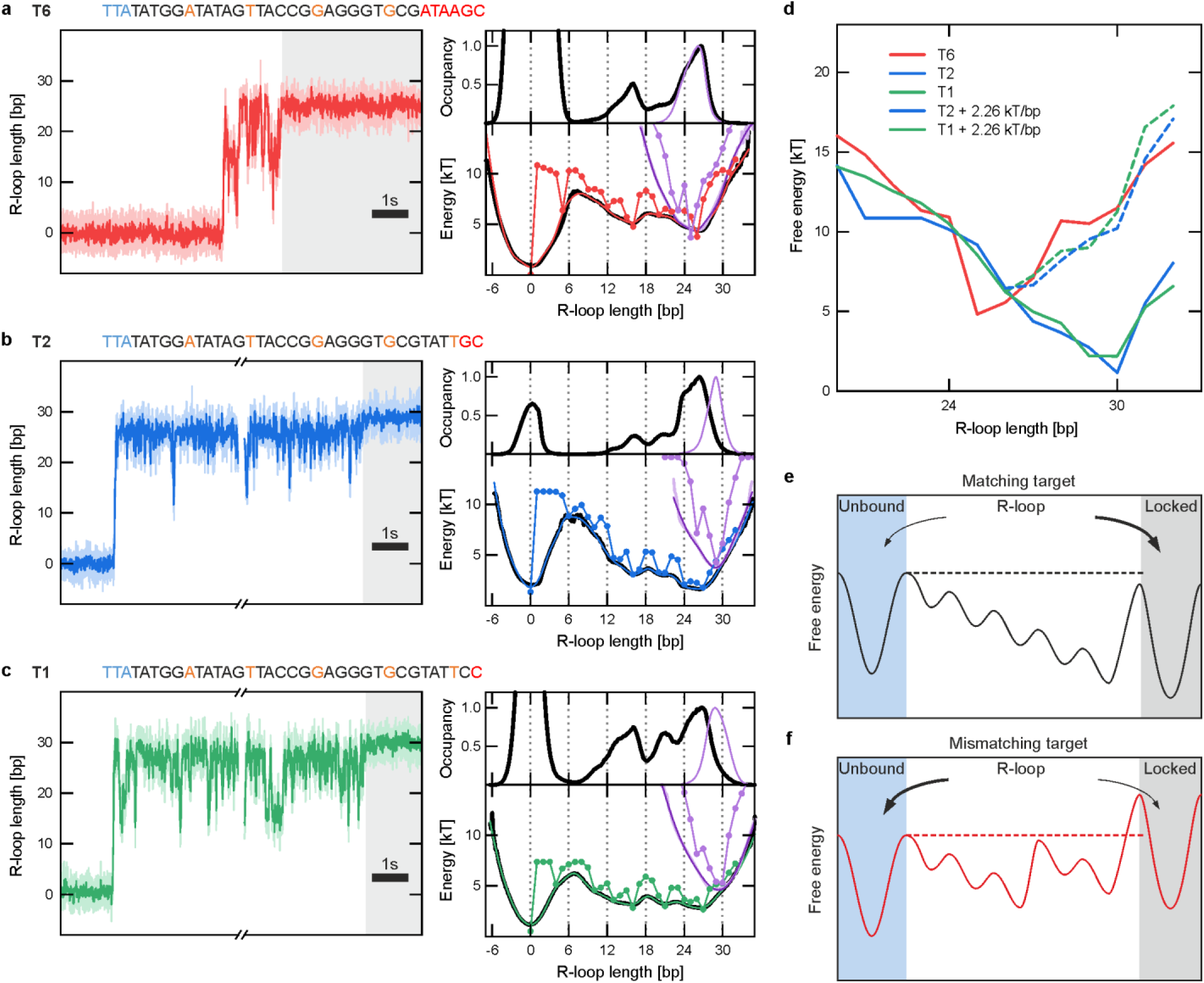
R-loop free energy landscapes before and after locking for the T6, T2 and T1 targets. **a-c,** (left) R-loop length trajectories including a locking event measured on targets with (**a**) six, (**b**) two and (**c**) one terminal mismatches (T6, T2, T1 targets). Data was taken at 3947 fps (light colours) and smoothed with a 100-point sliding average (dark colours). The extensive R-loop length fluctuations before locking became suddenly quenched upon irreversible locking indicating a tightly constrained R-loop length (grey shaded area). (Right) Histograms of the measured R-loop lengths as well as apparent (solid lines) and deconvolved (lines and circles) energy landscapes before (black, red, blue and green colours) and after locking (purple colour). For the T2 and T1 targets, a pronounced shift of the predominant R-loop length from 27 to 30 bp is observed upon locking. For the T6 target, the predominant R-loop length of ∼26 bp remains unchanged. **d,** Deconvolved free energy landscapes of the locked state for the T6, T2 and T1 targets (solid lines). As an attempt to reconstruct the free energy landscape of the T6 target from the landscapes of the other targets, we added a mismatch penalty of 2.26 *k_B_T* to the T2 and T1 landscapes for each additional mismatch that the T6 target comprises (dashed lines). The mismatch penalty agrees well with the average base-pairing energy of nucleic acid duplexes, suggesting that differences between the locked-state energy landscapes of the different targets and different equilibrium lengths arise from the additional mismatch penalties. Notably, larger R-loop lengths remain transiently accessible on the T6 target as indicated by the shallower slope of the landscape for lengths ≥ 26 bp). In contrast, smaller R-loop lengths appear to be transiently accessible on the T2 and T1 targets as indicated by the shallower slopes for lengths ≤ 30 bp. **e-f,** Scheme of the observed free energy landscapes of R-loop formation without (**e**) and with (**f**) an internal mismatch. Shaded areas indicate the unbound (blue) and the locked state (grey). In absence of a mismatch the terminal barrier is slightly lower than the initial barrier for PAM binding such that locked R-loop formation occurs at high probability. The penalty induced by the mismatch lifts the terminal barrier significantly up such that R-loop collapse becomes favoured over locked R-loop formation and the target is typically rejected. Such an energetics of R-loop formation supports a kinetic target discrimination mechanism^30^. Similar heights of terminal and initial barrier on a fully matching target ensure hereby highly efficient R-loop formation and high specificity at the same time

## SUPPLEMENTARY DISCUSSIONS

### S1 Frequency spectrum of the nanorotor fluctuations

The spatio-temporal resolution of twist measurements depends on the magnitude of the constrained nanorotor fluctuations when averaged over a given time window. Particularly, it is determined by the mean-square displacement of the angular fluctuations 〈φ^2^〉, as well as on their characteristic time scale given by the response time *τ* of this overdamped system. 〈φ^2^〉 is related to the torsional spring constant *κ* of the nanorotor according to 〈φ^2^〉 = *k_B_T*⁄*K*. The response time, which equals the time over which the angular fluctuations are correlated is given by τ = γ/*κ*, where *γ* is the rotational hydrodynamic drag coefficient of the nanorotor.

*κ* and *γ* of a particular nanorotor can be obtained by analysing the frequency spectrum of the angular fluctuations. The power spectrum of a simple harmonic oscillator in the limit of strong damping is a Lorentzian function given as^39^:

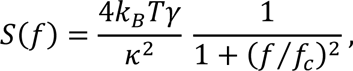

where *f_c_* = 1⁄2πτ = *K*⁄2πγ is the so-called cut-off frequency above which spectral contributions vanish. Below *f_c_*, the power spectrum is approximately constant in agreement with uncorrelated (white) noise. Averaging of the particle position over the integration time of the camera as well as aliasing change the measured signal, such that the actually observed power spectrum has the following form^40,41^:

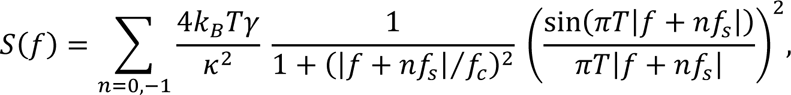

where *f_s_* is the sampling frequency of the camera and *T* the integration time, with *f_s_* = 1/*T* for all our measurements. Using this formula to fit the experimentally obtained power spectrum^40^ allowed us to determine *κ* and *γ* (Extended Data Fig. 6).

### S2 Spatio-temporal resolution of the nanorotor construct

The spatio-temporal resolution of a nanomechanical probe is proportional to the root-mean-squared (RMS) amplitude of the probe fluctuations σ_φ_(*t*) after the probe signal is filtered with a sliding average of width *t*. For a linear system with a single overdamped fluctuation mode, the time interval *t* contains approximately *N_uncorr_* ∼*t*/τ uncorrelated position segments (if *t* ≫ τ). The RMS amplitudes before and after filtering are thus related by:

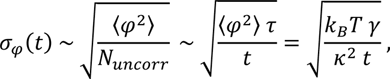

i.e. the spatio-temporal resolution can be increased by increasing the probe stiffness *κ* and/or decreasing the viscous drag coefficient *γ*.

One way to determine the spatio-temporal resolution is to calculate the Allan deviation σ_*m*_. This yields the RMS deviation of neighbouring time intervals of length *t* of a signal. For a trajectory recorded at a constant sampling frequency, the square of the Allan deviation (i.e. the Allan variance) can be calculated by^42^:

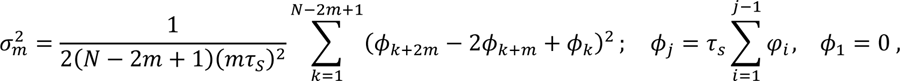

where τ_*S*_ = 1/*f_S_* is the sampling time, *t* = *m*τ_*S*_ the length of the time window and *N* the total number of data points. For a particle undergoing Brownian motion within a harmonic potential, the Allan deviation can be shown to take the form^42,43^:

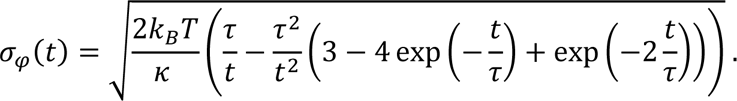

In agreement with our simplified derivation, it decays for *t* ≫ τ with 1/√ *t* until the thermal drift of the instrument limits the resolution on long time scales (Extended Data Fig. 6). For timescales similar to or shorter than the response time (*t* ≲ τ) it is limited by the slow response of the particle.

In order to determine the spatio-temporal resolution on short timescales, it is preferable to consider the RMS amplitude of the actual signal after filtering by a sliding average of width *t* which is given as^44^:

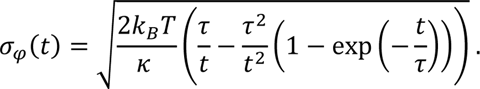

Again, it decreases for times *t* ≫ τ_*S*_ with 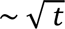. For small times, this relation approaches the total measured RMS before filtering.

To determine the temporal resolution over which a given change in angular position/untwisting can be resolved, we assumed that a signal change must exceed the probe fluctuation by a signal-to-noise ratio (SNR) of 3, and determined the corresponding time scale from the respective plots (Extended Data Fig. 6).

### S3 Design considerations and characteristics of the nanorotor

As discussed in section S2, the spatio-temporal resolution of a given mechanical probe can be improved by increasing the probe stiffness *κ* and/or decreasing the viscous drag coefficient *γ*. Both quantities are however subjected to technical as well as geometric constraints.

The drag coefficient of a rotating sphere is given by

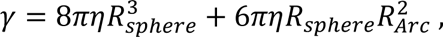

where *η* is the viscosity of the medium, *R_sphere_* the sphere radius and *R_Arc_* the distance of the sphere centre from the rotation axis. The first term in this equation is due to the rotation of the sphere around its centre and the second term due to the translational motion of the sphere centre on a circular path. *R_Arc_* needs to be larger than the largest lateral fluctuations of the DNA at the attachment point, otherwise the circular trajectory of the sphere could not be faithfully resolved. To achieve this criterion with a 5σ confidence interval for minimal forces of 2 pN and a height of the attachment point above the surface of ∼120 nm (see below), we chose *R_Arc_* ≈ 80 nm. For typical rotor bead assays^19^ the sphere would be directly attached to the DNA duplex, such that *R_sphere_* = *R_Arc_* and 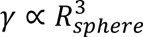. From the expression above, one sees, however, that the drag coefficient can be significantly decreased by reducing *R_sphere_* while keeping *R_Arc_* constant (eventually approximating a linear dependence on *R_sphere_*). This motivated the design and construction of the origami rotor arm.

To ensure detectable scattered light from the AuNP for tracking, we chose *R_sphere_* ≈ 25 *nm* (*R_sphere_* = 24 nm ± 4 nm, s. d., N = 328 as determined from TEM imaging) and constructed a rotor arm with *R_rotor_* = 56 nm length, nominally yielding *R_Arc_* = 81 nm. The mean experimentally determined arc radius, given by the major axis of the ellipse fits (see Methods) was 〈*R_Arc_* 〉 = 97 ± 10 nm. The difference can be attributed to two factors. The zipper like AuNP attachment geometry in combination with the steric hindrance between DNA coated AuNP surface and nanorotor adds an additional spacing of 2 to 3 nm. Additionally, steric hindrance and charge repulsion favour binding at pointed portions of the AuNP, such as the end of the long axis of an ellipsoidal AuNP or local protrusions of its surface (see Extended Data Fig. 1) where more space is available for neighbouring DNA strands to avoid each other. This locates the centre of the AuNP further from the dsDNA axis attachment point.

From the experimentally determined drag coefficients, we furthermore determined the mean apparent hydrodynamic radius of the AuNP using the equation from above. We observed *r*_AuNP_ = 30 nm ± 5 nm. The increased hydrodynamic radius compared to the radius from TEM measurements can in part be attributed to the functionalization of the nanoparticle, which adds a high density 20 nt poly-thymidine brush to its surface. Additionally, the contribution of the DNA rotor arm itself was neglected in the formula, which will increase the overall drag resulting in an overestimation of the actual hydrodynamic radius.

We chose a height of the AuNP centre above the surface of ∼118 nm, being more than four times the AuNP radius to avoid an increased viscous drag due to surface proximity^45^ as well as the sticking of AuNPs to the surface. To increase the torsional rigidity, the amount of linear dsDNA positioned below the AuNP was kept at a minimum. It comprises the Cascade target sequence and the dsDNA-origami interface (total length of 80 bp, corresponding to *L_DNA_* = 27 nm). The remaining part was a six-helical stem provided by the 105 nm origami rotor arm structure of which *L_rotor_* = 91 nm were located below the centre of the attachment points of the AuNP. Such a six-helical stem has a ∼6-fold increased torsional persistence length (*p_rotor_* = 530 nm) compared to dsDNA (*p_DNA_* = 92 nm)^46^. The torsional stiffness of the full nanorotor tether is given as a linear combination of two torsional springs:

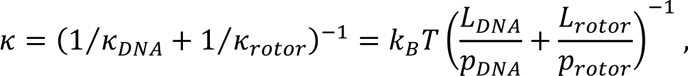

using *κ* = *k_B_T p*⁄*L* with *p* being the torsional persistence length of the given tether. For the given geometry, the dsDNA and the nanorotor part both provide significant contributions (despite very different lengths) resulting in an expected total spring constant of *κ* = 8.9 pN nm⁄rad. Experimentally we observed *κ* = 5.2 ± 0.2 pN nm⁄rad. The deviation is attributed to an increased length of the dsDNA part. The surface attachment handle contains only a limited number (∼30) of digoxigenin anchors at random positions, such that always a part of the handle before the first digoxigenin attachment adds to the dsDNA tether length.

Experimental values for the mean torsional stiffness, the drag coefficient and the time required to reach a spatial resolution of a single base pair at a SNR of 3 are presented and compared to previous studies in Extended Data Table 1. Compared to DNA-attached rotor beads used by Ivanov et al. in actual measurements of R-loop formation, an improvement of the spatio-temporal resolution of almost one order of magnitude was achieved. This was due to the increased torsional stiffness of our nanorotor which reduced the overall RMS of the fluctuations and the response time. While the approach of Kosuri et al. yielded an even higher spatio-temporal resolution, the use of organic fluorophores limited the total measurement time to 3-4 minutes (compared to up to 80 minutes in this study) which would not have been sufficient to sample the entire energy landscape with sufficient statistics particularly at elevated energy values.

### S4 Torque measurements

The nanorotor construct could also be employed to measure torque within the dsDNA spacer above the nanorotor (Extended Data Fig. 7). For torque measurements upon DNA twisting, we first identified functional molecules that were torsionally constrained towards both sides of the nanorotor, i.e. towards the surface as well as towards the magnetic bead. To this end, we twisted the molecules and verified the occurrence of the typical DNA length reduction due to the formation of writhe in form of a plectonemic superhelix (see also below)^47^.

Generally, the torque can be calculated from the angular deflections of the nanorotor with respect to the surface (φ_*rotor*_) as well as with respect to the magnetic bead (φ_*bead*_ − φ_*rotor*_) as:

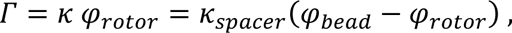

where φ_*bead*_ denotes the angular displacement applied to the magnetic bead and *κ* as well as *κ_spacer_* are the torsional spring constants of the elements below and above the nanorotor arm. For calibrating either spring constant (close to zero torque) using power spectral density analysis (see above), one has to consider that a nanorotor displacement acts in parallel against both springs, such that calibration yields the total spring constant *K_tot_* = *K* + *K_spacer_*. Transforming the torque relation yields: *κ_spacer_*⁄*K* = φ_*AuNP*_⁄(φ_*bead*_ − φ_*rotor*_). Combining the two expressions gives:

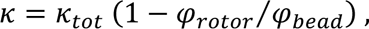

φ_*rotor*_⁄φ_*bead*_ is obtained in a twisting experiment for extended (non-buckled) DNA from the slope of the nanorotor displacement vs. the applied turns of the magnetic bead (Extended Data Fig. 7).

When carrying out nanorotor-based torque measurements during DNA twisting, the expected characteristic behaviour was observed^24,36,48^. Upon moderate positive or negative DNA supercoiling (i.e. overwinding or underwinding of the helix, respectively), the DNA length remained initially constant, while the torque changed linearly with the applied turns (Extended Data Fig. 7b). In this regime, the torque in the spacer DNA is given as:

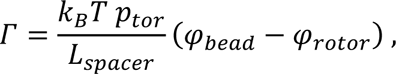

with *L_spacer_* being the contour length of the DNA spacer and Τ = *K* φ_*rotor*_ the acting torque. From fitting the experimental torque relationship with this relation, a torsional persistence length of the dsDNA spacer of *p_tor_* = 102 ± 3 nm was obtained in good agreement with previous measurements^36^. The linear torque regime was abandoned once critical torques were reached. For positive supercoiling and moderate force an abrupt length decrease was observed at which the molecule buckled^24,36,48^, which was accompanied by a moderate torque ‘overshoot’. Subsequently, the molecule length decreased linearly with the applied turns due to the growth of a plectonemic superhelix during which the torque remained constant (Extended Data Fig. 7b). In agreement with previous measurements, the torque plateau increased with increasing force. At a force of 5.7 pN a torque plateau was observed despite the absence of buckling. In this case, the applied twist is released by p-DNA formation^25^ which is an overwound DNA structure. For negative supercoiling^36^, a force-independent torque plateau was reached at 10 ± 1 pN nm at which excess twist is accommodated by local melting or Z-DNA formation^19^. Occasionally, transient melting in the dsDNA region below the nanorotor was also seen, providing short lived, strong negative angular deflections (Extended Data Fig. 7b). Overall, we were able to quantitatively monitor the torque for the entire spectrum of DNA twisting transitions. We note that our torque measurements were performed in less than three minutes of measurement time per force (0.5 turns/s) due to the high torque resolution of the nanorotor design of 1 pN nm per 38 ms.

### S5 Deconvolution of the Energy Landscape of R-loop formation

The high spatio-temporal resolution of the nanorotor-based DNA untwisting measurements allowed to follow the dynamic sampling of the different R-loop lengths. From the measured distribution of the angular nanorotor positions *P*^′^(φ_*rotor*_), an apparent energy landscape *G*^′^(φ_*rotor*_) = −*k_B_T* ln[*P*^′^(φ_*rotor*_)] was calculated using the Boltzmann distribution (Fig. 2, Extended Data Fig. 8). To obtain the actual energy landscape of the forming R-loop, one has to consider that the DNA untwisting by the R-loop is overlaid by thermal fluctuations of the nanorotor that are also seen in absence of an R-loop (see main text, Fig. 1d, top). This broadened the measured angular distributions and thus the apparent energy landscapes. Mathematically, this corresponds in first approximation (neglecting coupling effects) to a convolution of the real distribution of DNA untwisting by the R-loop *P*(φ) with the nanorotor position distribution *K*(φ) as a kernel:

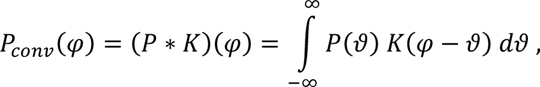

where *K*(φ) can be directly measured in the absence of Cascade. The Kernel generally followed a distribution that approximately resembled a Gaussian distribution, with a typical standard deviation of less than two base pairs (Fig. 2b, Extended Data Fig. 8). A more precise approximation of *K*(φ) was achieved using a superposition of three Gaussians of the same width where a dominant peak was flanked by two minor peaks at a distance of about 1 rad. The deviation from a purely Gaussian profile could be caused by fluctuations of the AuNP around its attachment to the rotor arm as well as transient opening and closing of the first base pairs at the nanorotor – dsDNA interface.

The discrete nature of the R-loop base-pairing motivated us to use a discrete free energy landscape *G*(*n*) and length distribution *P*(*n*) for the R-loop that are related by the Boltzmann distribution according to:

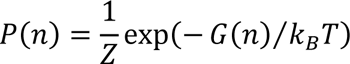

where the partition function 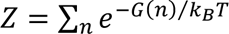 ensures the normalization of *P*(*n*). With Δφ_bp_ being the untwisting angle per base pair, the DNA untwisting of an R-loop of length *n* can be expressed as:

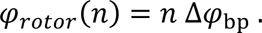

The convolution simplifies then to a sum:

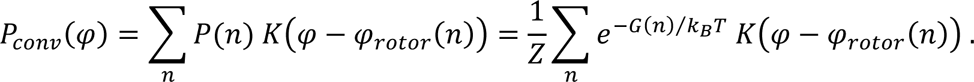

To determine the actual free energy landscape of R-loop formation *G(n)*, we performed weighted least-squares fits of the resulting convolved energy landscape *G_conv_*(φ) = − ln(*P_conv_*(φ)) to the measured apparent energy landscape *G*^′^(φ_*rotor*_). Weights *w*(φ) were set to 0 for values less than −2 bp as well as positions larger than 34 bp. To avoid boundary effects from a low statistics, a weight of 0.2 was used for positions between −2 and 8.2 bp as well as 32 and 34 bp, while for the remaining positions from 8.2 to 32 bp of R-loop length a weight of 1 was applied.

In a first round, a global fit was performed to the measured apparent free energy landscapes of a given construct (*N* = 4 to 5). Free fit parameters were the values of the ‘real’ R-loop length distributions *P*(*n*) for R-loop lengths *n* from 0 to 32, the single base pair untwisting angle Δφ_bp_ and the angular offset φ_0_, which defines the mean angular position of the nanorotor in absence of Cascade. After finding a consistent best-fit value of the single base pair untwisting angle of Δφ_bp_ = 0.573 rad = 32.8°, Δφ_bp_ was subsequently taken as a fixed parameter to allow a more rigorous fitting at higher computation efficiency. In a second round we performed individual fits to the individual free energy landscapes of the different characterized molecules, where only the values of the R-loop length distributions *P*(*n*) were used as parameters. To deconvolve the energy landscape of the locked state, a separate set of individual fits was performed. In this case the weight was derived from the apparent length distribution as: *w*(φ) = *P*′(φ)^0.25^, while the offset parameter was taken from the global fits of the free energy landscapes.

A deconvolution based solely on a least-mean-square fit resulted in unrealistically high energy barriers between local energy minima as an attempt to compensate for the statistical noise in the energy landscapes (see result for *b* = 0 in Extended Data Fig. 9). To avoid this, we applied a maximum entropy deconvolution algorithm which additionally tries to minimize statistical bias, i.e. large local alterations in the landscapes. To calculate the statistical bias Δ*G_entr_*(*n*) of an energy landscape *G*(*n*) during fitting, we constructed a reference landscape 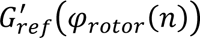 based on the measured apparent energy landscape *G*^′^(φ_*rotor*_(*n*)) between 7 and 32 bp of DNA untwisting. Since the dominant unbound state distorted the energy landscapes between 1 and 7 bp, we extrapolated 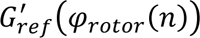 in this range from a linear fit to *G*^′^(φ_*rotor*_(*n*)) between the energy maximum at ∼7 bp and the minimum at 26 bp (see dashed blue line in Extended Data Fig. 9a). For the target containing the single mismatch (T6-M17) we additionally added to the energy values of all positions ≥ 17 bp the mismatch penalty of 4.5 *k_B_T*) which we determined from the difference between the apparent energy landscapes of the mismatch and the T6 target (see Extended Data Fig. 10b). The statistical bias Δ*G_entr_*(*n*) was then obtained as:

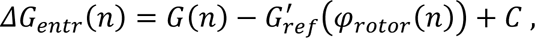

where *C* = 5 was an empirically chosen offset ensuring that all values were positive. Δ*G_entr_*(*n*) was normalized to yield 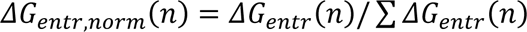. Next, the Shannon entropy for the normalized energy bias was calculated:

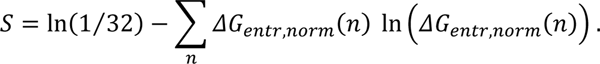

*S* approaches zero if *G*(*n*) equals the measured apparent energy landscape *G*^′^(φ_*rotor*_(*n*)) and yields increasingly negative values for increasing difference between the landscapes. The calculated entropy was added as a penalty to the least square residue of the fit:

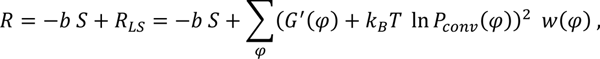

where *R* is the total residue, *R_LS_* the pure least square residue between the convolved energy landscape and the apparent energy landscape from the measurement, *w*(φ) the applied weights as described above and *b* the entropy scaling factor. Following total residue minimization, increasing *b* provided deconvolved energy landscapes *G*(*n*) with decreasing energy barriers, (see Extended Data Fig. 9a). We performed the maximum entropy deconvolution across a range of entropy scaling factors. The extracted deconvolved energy landscapes were subsequently applied in Brownian dynamics simulations to investigate which set of parameters best described the measured dynamics of the R-loop length fluctuations at different timescales.

### S6 Brownian Dynamics Simulations of R-loop dynamics

To model the dynamics of our measurements of R-loop formation, we approximated the setup as a coupled system of the nanorotor undergoing one-dimensional diffusion in a harmonic potential and of the R-loop undergoing a random walk in discrete steps within the R-loop energy landscape.

The Brownian dynamics simulations were performed in incremental time steps of Δ*t* = 10 µs. During each step, the nanorotor was moved by a random diffusive displacement Δφ_*diff*_ which was drawn from a normalized Gaussian distribution with variance 〈Δφ_*diff*_^2^〉 = 2*D*Δ*t* with *D* = *k_B_T*⁄γ being the rotational diffusion coefficient. Additionally, we considered the back-driving torque on the nanorotor Γ = −*k_tor_*ϑ = −*k_tor_*[φ − φ_*R*_(*n*)] due to angular displacements ϑ of the nanorotor from the equilibrium position φ_*R*_(*n*) given by the current R-loop length. In the limit of strong damping, the back-driving torque causes an angular drift velocity ω = Γ⁄γ. The total nanorotor displacement per time increment Δ*t* was then derived as:

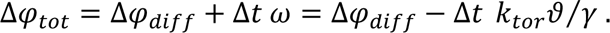

Parallel to the simulation of nanorotor displacements, we let the R-loop elongate or shorten in single base pair increments. The corresponding probabilities *p*_−_(*n*) and *p*_+_(*n*) for elongation and shortening per time increment Δ*t* were for sufficiently short increments given by the local backward and forward stepping rate constants:

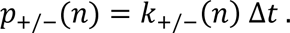

For an unbiased energy landscape, the local forward and backward stepping rate constants are equal and we furthermore considered them to be position independent, i.e. described by a single rate constant *k_step_*. In case of bias between two neighbouring positions *n* and *n* + 1, the ratio between the corresponding forward and backward rate constants is related to the bias according to the principle of detailed balance:

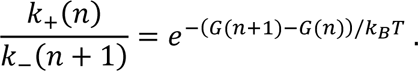

We assumed that the activation barrier for the transition is centred between the two positions. In this case the activation barrier is changed by the bias by (*G*(*n* + 1) − *G*(*n*))/2 such that we get according to the Arrhenius law for both rate constants:

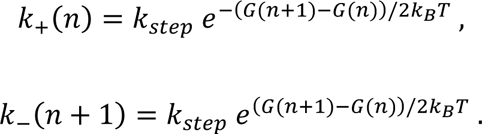

In addition to the statical bias from the R-loop energy landscape, the R-loop formation experiences additional dynamically changing bias due to the torque from the angular nanorotor fluctuations. Positive displacements ϑ from the equilibrium position φ_*R*_(*n*) (i.e. towards DNA unwinding) will assist R-loop elongation, while negative displacements ϑ will assist R-loop shortening. To determine how the torque changes the height of the activation barriers, we calculated for a fixed nanorotor position ϑ the change in torsional energy (*G_tor_* = *k_tor_*ϑ^2^/2) between the equilibrium position φ_*R*_(*n*) of the R-loop and the transition state located at ± Δφ_bp_/2 from φ_*R*_(*n*). For R-loop elongation/shortening we get then the following changes of the activation energy:

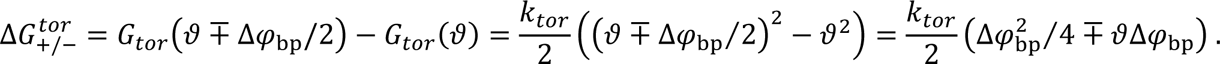

This changes the stepping rate constants as function of the nanorotor position to:

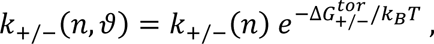

which provides an additional mutual coupling between the R-loop and nanorotor dynamics beyond the centring of the nanorotor position around φ_*R*_(*n*).

Each simulation was run for a total simulated time of 10 min. For further analysis, trajectories of R-loop length and nanorotor position were averaged over successive time intervals of 250 µs and correspondingly downsampled. The simulations were conducted for a range of different stepping rates *k_step_* and for different deconvolved energy landscapes produced by the different entropy scaling factors *b* (see deconvolution chapter).

### S7 Comparing measured and simulated R-loop dynamics using transition maps

In order to find the stepping rate *k_step_* and the deconvolved energy landscapes that best describe the full R-loop dynamics, we used a discrete set of transition maps to compare the time evolution of measured and simulated R-loop dynamics (see Fig. 3 and Extended Data Fig. 9).

A single transition map showed for R-loop lengths >7 bp the probability that an apparent R-loop length from the nanorotor measurements *n*(*t*) (x-axis) changes after a time Δ*t* into an R-loop length *n*(*t* + Δt) (y-axis). For each initial R-loop length, the probabilities were normalized, to reveal the fate of an R-loop of a given length. Transition maps of the experimental data were created for time intervals Δt of 2, 5, 10 and 25 ms. The measured R-loop length distributions in the transition plots diffusively broadened over time. For 2 ms and, to a lesser extent, also 5 ms, the presence of four pronounced spots in the transition plots indicated that the R-loop remained confined within a given energy well (Extended Data Fig. 9b) formed by the energy barriers at R-loop lengths of 6, 12, 18 and 24 bp (Extended Data Fig. 9a). This corresponds to intra-well diffusion about the well centres at 9, 15, 21 and 26 bp. When looking at the transition for intervals of ≥ 10 *ms*, significant inter-well transitions were observed, i.e. the R-loops frequently left the initial well such that the initial sharp dots of the wells became fuzzy (Extended Data Fig. 9b).

We next calculated transition plots for the simulated R-loop dynamics across the range of tested stepping rates *k_step_* and energy landscapes (as determined by the entropy scaling factor *b*). To find the parameter set that best described the measured dynamics, we calculated for each parameter set the residue over all time intervals between the experimental and the simulated transition plots for R-loop lengths between 15 and 26 bp. Minimization of the residue score provided the best fit parameter set (Extended Data Fig. 9c). The deconvolved energy landscape associated with the best-fit value of *b* was taken as “best-fit” energy landscape. Averaged energy landscapes from at least 5 molecules were used to display the sequence-specific energy landscapes in Fig. 3c, main text.

### S8 Estimating inter-well transition rates

Based on the calculated transition maps (Fig. 3 and Extended Data Fig. 9) and the MSD analysis (Extended Data Fig. 11) we concluded that the R-loop dynamics is governed on short time scales by diffusion within the local energy wells (intra-well diffusion) and on longer time scales by inter-well transitions. To estimate the inter-well transition rate constants, we defined successive wells of 6-bp width centred on R-loop lengths of 9 bp, 15 bp, 21 bp and 26 bp. For any time point where the apparent R-loop length was found at a well centre, we determined the time required to escape the initial well by reaching the centre of either adjacent well. For each well, total escape rate constants *k_i,esc_* were determined by fitting the survival distribution of the escapes with an exponential function. The rate constants for forward (*k_i,+_*) and backward (*k_i,−_*) escapes from a particular well *i* were obtained by multiplying *k_i,esc_* with the escape probabilities for the given direction:

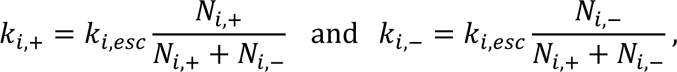

where *N_i,+_* and *N_i,−_* indicate the number of observed escapes from well *i* in forward and backward direction, respectively.

From the reciprocal of the escape rates the average dwell time per well can be obtained: *t_i,esc_* = 1⁄*k_i,esc_* (4.9, 29, 9.4 and 39 ms for wells 2,3,4 and 5 of the T6 target respectively, see also Extended Data Fig. 11). The measured escape rates can also be used as stepping rates when describing diffusive R-loop formation as a 6-bp random walk. The diffusion constant for such a random walk would then be given as:

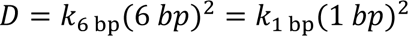

with the right-hand side of the equation describing a corresponding random walk with 1-bp step size along a flat energy landscape and the same diffusion constant. Using 〈*k_esc_* 〉 = 21 ms then yields a corresponding 1-bp stepping rate of *k*_1 bp_ = 1700 s^−1^ in good agreement with previous modelling of the R-loop formation dynamics^16^.

## SUPPLEMENTARY METHODS

### Purification of Cascade complex

*Streptococcus thermophilus DGCC7710* Cascade complex was heterologously expressed in *E. coli* BL21 (DE3) cells using pACYC pCRh encoding homogeneous CRISPR region^49^. The Cascade complex was purified as described^2^.

### Assembly of the nanorotor construct

The nanorotor construct was prepared from 5 different parts (Extended Data Fig. 3): A ∼620-bp digoxigenin-modified surface attachment handle, a 47 bp target sequence, the origami rotor arm, a 7.5-kbp dsDNA spacer and a ∼620-bp biotin-modified attachment handle. All components contained suitable sticky ends to allow assembly of the full nanorotor in a final ligation step.

For the preparation of surface and bead attachment handles, 1.2-kbp long PCR fragments were produced using Taq DNA Polymerase (NEB) and pBlueScript II SK (+) as a template, such that the multiple cloning sites of the vector were located in the fragment centre (See Extended Data Table 3 for primers). Multiple biotin or digoxigenin modifications were introduced in the fragments, by supplementing the PCR reactions with either Biotin-16-dUTP (Jena Bioscience) or DIG-11-dUTP (Jena Bioscience) at a 1:10 ratio to dTTP. The biotinylated and digoxigenin-modified fragments were purified (Clean & Concentrator-25, ZymoResearch) and digested with SpeI and NotI (both New England Biolabs) to produce ∼620-bp modified fragments with a single sticky end.

The 47-bp DNA duplexes containing the Cascade target sequences were hybridized from two synthetic oligonucleotides (Extended Data Table 2) in 10 mM TRIS-HCl (pH 8.0), 200 mM NaCl, 5 mM MgCl_2_ and 1 mM EDTA by heating to 95°C followed by slow cooling to room temperature at −1 K/min.

The DNA origami rotor arm was designed using CaDNAno^18^ (Extended Data Fig. 1, 2). For its assembly, 10 nM 8064 nt long single-stranded scaffold DNA (p8064 variant of the M13 bacteriophage genome, MWG Eurofins, Ebersberg, Germany) was mixed with the 206 DNA staples (75 nM each) in folding buffer (11 mM MgCl2, 5 mM TRIS-HCl (pH 8.0), 1 mM EDTA). Subsequently, the solution was heated to 95°C and slowly cooled to room temperature over 15 h as described^46^. The origami nanostructures were then purified twice, using PEG precipitation^50^ to remove excess staples. The nanostructure concentration was determined from UV absorbance measurements at 260 nm. Subsequently, the secondary interface oligomers (Extended Data Fig.3c) were added at a 10-fold molar excess over origami structures and annealed to primary interface overhangs by heating to 40°C and subsequent slow cooling to room temperature at −0.2 K/min. The rotor arms were then again purified from excess oligonucleotides using PEG precipitation.

The 7.5-kbp dsDNA spacer was made via PCR using Q5 polymerase and lambda DNA as template. Each primer (Extended Data Table 3) contained a site for a restriction enzyme, such that sticky ends were produced at both termini by digestion with Nb.BbvCI and SpeI.

For the ligation of the full nanorotor DNA construct, the five different segments were mixed according to the amounts given in Extended Data Table 4. The ligation was then carried out in a total volume of 20 µl for 1 h at room temperature using 400 units of T4 Ligase (NEB). Excess handles and target duplexes were subsequently removed using PEG precipitation and the sample was resuspended in nanorotor buffer containing 11 mM MgCl_2_, 5 mM TRIS-HCl pH 8.0 and 150 mM NaCl (Extended Data Fig. 3).

### Preparation of TEM samples

For TEM imaging, a 5 μl droplet containing 1-2 nM of DNA origami nanorotors was placed onto a glow-discharged, carbon-coated TEM grid for 5 min, followed by staining with a filtered solution containing 2% uranyl formate and 5 mM NaOH. TEM imaging was performed on a Jeol JEM2100Plus transmission electron microscope at 200 kV.

### Preparation of magnetic tweezers experiments

For flow cell preparation, two 60 mm x 24 mm cover slides (Menzel, ThermoScientific) were sequentially sonicated in ultrapure water, acetone and isopropanol for 10 minutes each. After rinsing with ultrapure water, they were sonicated for an additional 20 minutes in 5 M KOH followed by another rinsing step with ultrapure water and drying with pressurized N_2_ gas. The top slide contained two holes of 2 mm diameter as inlet and outlet. The bottom slide was coated with polystyrene by spin coating using a 1% w/v polystyrene solution in toluene and 6000 rpm. Subsequently, the polystyrene layer was baked for 1 h at 150°C. Flow cells were assembled on a hot plate at 120°C as a sandwich of the two cover slides and a Parafilm® spacer in which the flow cell cavity was cut out. For surface functionalization, the flow cell was incubated over 24 h with 50 µg/ml solution of anti-digoxigenin (Bio-Rad) in phosphate buffered saline (PBS). It was then incubated over night with a 20 mg/ml solution of BSA (NEB) in PBS to prevent non-specific binding. Next, the flow cell was mounted into the holder of the microscope, connected to a pump and carefully washed with 1 M NaCl. 3 µm-sized carboxylated polystyrene beads (Invitrogen, Carlsbad, CA) serving as reference particles during bead tracking were added in 1 M NaCl and incubated over night to allow their firm adsorption on the bottom slide.

The 50 nm gold nanoparticles (AuNPs) of the nanorotors were synthesized as previously described and functionalized with 20 nt 3’-thiol-modified poly-thymidine oligonucleotides using salt-aging^51,52^. DNA-coated AuNPs were mixed in a 5-fold molar excess with nanorotor DNA constructs in nanorotor buffer, heated to 40°C and cooled to room temperature at −0.2 K/h.

For magnetic tweezers experiments, 2 µl magnetic beads of 1.05 µm diameter (MyOne, Invitrogen) were washed twice in PBS. They were then incubated with 0.2 fmol AuNP-nanorotor constructs for 5 minutes at room temperature in 10 µl preparation buffer (10 mM TRIS-HCl at pH 8.0, 150 mM NaCl, 6 mM MgCl_2_, 0.01% Pluronic F-127 and 0.1% Tween). Afterwards, preparation buffer was added up to 100 µl, the magnetic particles were pelleted and the supernatant including excess AuNPs and ligation side products was removed. The beads with bound nanorotor constructs were then resuspended in 100 µl of preparation buffer.

Directly before a measurement, preparation buffer was flushed into the flow cell and left for 10 min of incubation. The buffer was then replaced by flushing 300 µl nanorotor buffer (see above). 50 µl magnetic beads with bound nanorotor constructs were added and incubated for 5 minutes. Unbound beads were then removed by adding another 300 µl of nanorotor buffer supplemented with 8 µg/ml catalase (Sigma-Aldrich/Merck) and 20 µg/ml glucose oxidase (Sigma-Aldrich/Merck) to prevent damage of the DNA during laser illumination^53^. This buffer was replaced every 60 minutes, if required.

### Ultrafast twist measurements

Measurements were performed in a setup combining magnetic tweezers with scattered light microscopy (Fig. 1c, Extended Data Fig.4). Force onto nanorotor-tethered magnetic beads was generated using the strong gradient of a magnet pair (W-05-N50-G, Supermagnete) mounted in a magnetic holder made of iron, which could be translated and rotated using computer-controlled stages (M-122.2DD and C-150.PD, Physik Instrumente). For real-time measurements of the DNA length, the positions of the magnetic bead and a non-magnetic reference bead (providing drift-elimination, Extended Data Fig. 5c) were tracked simultaneously by imaging each bead within a 160 px x 160 px area at 1000 fps using an EoSens CL MC-1362 CMOS camera (Mikrotron, Germany) and applying GPU-assisted real-time particle tracking^20^. The scattered light from the AuNP of a given nanorotor was imaged within a 32 px x 32 px area at 3947 frames per second (Extended Data Fig. 5b) using an ORCA-Flash4.0 V2 sCMOS camera (Hamamatsu, Japan). The signals of the two cameras were correlated in time with sub-millisecond resolution as verified by monitoring stage induced displacements of surface adhered reference beads and AuNPs in their respective channel. Suitable nanorotor constructs were identified by searching for a magnetic bead-tethered DNA construct of the correct length and ensuring the presence of a mobile, yet nanorotor-constrained AuNP (See Fig. 1d). Magnet rotations were applied to supercoil the spacer DNA in direct torque measurements (Extended Data Fig. 7). They were also used to select constructs that were torsionally unconstrained towards the magnetic bead in Cascade measurements.

To determine the angular position of a nanorotor in time, the intensity profile of the AuNP was fitted to a 2D Gaussian within a 13 px x 13 px area centred around the pixel with maximum intensity (Extended Data Fig. 5c) yielding the AuNP position. In order to reduce the influence of background noise, the fit used the square root of the intensity as weight if above a chosen threshold or a zero weight if below the threshold. As threshold, a fifth of the intensity of the 95% percentile over the fitting area was taken. Thermal drift of the setup was removed by subtracting the reference bead position. The influence of the pendulum-like fluctuations of the bead-tethered DNA onto the AuNP position was corrected using the position of the magnetic bead (Extended Data Fig. 5c). The corrected XY positions of the AuNP were found to follow a slightly eccentric ellipse. This was attributed to a slightly tilted rotation plane of the nanorotor (Extended Data Fig. 5d). To obtain correct nanorotor angles, the AuNP positions along the minor ellipse axis^54^ were increased to yield a circular trajectory. The nanorotor angle was then provided by the polar angle of the AuNP position with respect to the circle centre.

For twist and torque measurements, molecules that had an expected end-to-end distance of around 2.9 µm and were equipped with a fluctuating AuNP, were selected. For direct torque measurements, supercoilable molecules for which a DNA shortening upon twisting could be seen were selected, while for R-loop formation experiments non-supercoilable molecules were chosen. In both cases the nanorotor had to exhibit constrained rotational fluctuations. Power spectral density analysis of the angular trajectories of the nanorotor yielded the torsional spring constant and the hydrodynamic drag coefficient of the particular nanorotor (Extended Data Fig. 6). The drag coefficient was used to obtain an apparent hydrodynamic radius for the AuNP considering the geometry of the nanorotor. Calculation of the Allan deviation as function of averaging time was used to estimate the spatio-temporal resolution of the nanorotor (Extended Data Fig. 6). More detail on the analysis of the measurements is provided in the Supplementary Discussions. Measurements in absence of protein were conducted for at least 45 s for each molecule.

Single-molecule torque measurements were conducted by rotating the magnet pair to supercoil the tethered DNA construct and simultaneously measure the angular displacement of the nanorotor. Supercoils were added at a rate of one turn per second in a range from −35 to 65 turns. Using the calibrated torsional stiffness *κ* of the nanorotor allowed calculating the acting torque on the molecule as a function of the nanorotor deflection as: Τ = *K* φ_*rotor*_. The apparent torsional stiffness of DNA was obtained from the slope of the torque-turn curve in the linear regime (see Supplementary Discussions S4). This analysis confirmed the previously measured torsional properties of DNA.

### Real-time measurements of the dynamics of R-loop formation

After selecting and mechanically characterizing a suitable nanorotor, 2.5 nM *S. thermophilus* Cascade^2^ in measurement buffer was added to the flow cell. Ultrafast twist measurements were conducted until the nanorotor became dysfunctional, e.g. due to nicking of the dsDNA below the nanorotor by photo damage, AuNP detachment (photodamage at the AuNP attachments) or R-loop locking despite the present mismatches. Trajectories were recorded continuously for up to 4 minutes, to keep file sizes manageable. The total measurement time of subsequently recorded trajectories in presence of Cascade was typically tens of minutes, with a maximum of 80 min. From the recorded images of the AuNP, trajectories of the angular position of the nanorotor were calculated (Extended Data Fig. 5).

### Calculation of the energy landscape of R-loop formation

From the recoded trajectories of a given experiment a histogram of the angular positions of the nanorotor was calculated (See Supplementary Discussions S5). This directly allowed to determine the apparent energy landscape since the free energy *G*(φ) and the probability of encountering a given DNA untwisting *P*(φ) by the R-loop are related by the Boltzmann distribution *P*(φ)∼ exp(−*G*(φ)/*k_B_T*). The DNA untwisting angle by the R-loop and nanorotor angle are not identical due to the angular fluctuations of the nanorotor on top of the DNA untwisting. Thus, the probability distribution of the nanorotor angles *P*^′^(φ) allows only to calculate an apparent energy landscape *G*^′^(φ) = −*k_B_T* ln[*P*^′^(φ)] that is convolved with the approximate harmonic potential governing the nanorotor fluctuations. We determined the angular trapping potential of the nanorotor from enzyme-free measurements and used it as kernel for a subsequent maximum entropy deconvolution^55^. To obtain an estimate of the real energy landscape of R-loop formation *G*(φ) near base pair resolution, we deconvolved the apparent landscape *G*^′^(φ) for a set of entropy scaling factors and single base pair stepping rates *k_step_*. We next selected the parameter set that could best describe the experimentally observed dynamics of R-loop formation (see Supplementary Discussions S6). To this end we applied the deconvolved energy landscapes and stepping rates in Brownian dynamics simulations that coupled R-loop formation and nanorotor fluctuations and compared them with the experimental data using transition plots for a set of selected times (see Supplementary Discussions S7). For the T6-M17 target, we compared the mismatch penalty to the penalty of introducing a mismatch to a DNA duplex (see main text), calculated with NUPACK^56^.

